# Acute perturbation of retromer and ESCPE-1 leads to functionally distinct and temporally resolved defects in endosomal cargo sorting

**DOI:** 10.1101/836460

**Authors:** Ashley J. Evans, James L. Daly, Anis N. K. Anuar, Boris Simonetti, Peter J. Cullen

**Author notes:** Correspondence: Peter J. Cullen – and Ashley J. Evans.

## Abstract

The human retromer is a stable heterotrimer of VPS35, VPS29 and VPS26 whose principal role is to orchestrate the endosomal retrieval of hundreds of internalised cargo and promote their recycling to the cell surface; a prototypical cargo being the glucose transporter GLUT1. Retromer’s role in a distinct endosomal retrieval pathway, the retrograde sorting of the cation-independent mannose 6-phosphate receptor (CI-MPR) to the *trans*-Golgi network (TGN), remains controversial. Here we have developed and applied knocksideways to acutely inactivate retromer and by visualising the sorting of endogenous GLUT1 and CI-MPR provide insight into the temporal dynamics of endosomal cargo sorting in HeLa and H4 human neuroglioma cells. While retromer knocksideways led to the development of time-resolved defects in cell surface sorting of GLUT1 we failed to observe defects in the sorting of the CI-MPR. In contrast knocksideways of ESCPE-1, a key regulator of retrograde CI-MPR sorting, resulted in a time-resolved defect in CI-MPR sorting. Together these data provide independent evidence consistent with a comparatively limited role for retromer in ESCPE-1 dependent CI-MPR retrograde sorting in HeLa and H4 human neuroglioma cells.

## INTRODUCTION

The endosomal pathway functions as a major intracellular hub for the sorting of numerous integral proteins, which include signalling receptors, adhesion molecules, nutrient transporters, ion channels, and their associated proteins and lipids (collectively termed ‘cargoes’) (Maxfield and McGraw, 2004; Grant and Donaldson, 2009; Cullen and Steinberg, 2018). On entering the pathway, cargoes are sorted between two fates: they are either selected for degradation within the lysosome or retrieved from this fate and promoted for recycling to the plasma membrane and the *trans*-Golgi network (TGN) (Cullen and Steinberg, 2018). The efficient sorting of cargo is essential for normal cellular homeostasis and defects in sorting are increasingly linked with human physiology and pathophysiology (Schreij et al., 2016; Cullen and Steinberg, 2018).

Sequence-dependent cargo sorting for retrieval and recycling is orchestrated by highly conserved multi-protein complexes that include the retromer and retriever complexes, the COMMD/CCDC22/CCDC93 (CCC) complex, and the ‘*Endosomal SNX-BAR sorting complex for promoting exit 1*’ (ESCPE-1) complex (Seaman et al., 1998; Carlton et al., 2004; Phillips-Krawczak et al., 2015; McNally et al., 2017; Simonetti et al., 2019). These bind to sorting motifs present within the intracellular cytoplasmic domains of cargo either directly (Fjorback et al., 2012; Phillips-Krawczak et al., 2015; Bartuzi et al., 2016; Lucas et al., 2016; Kvainickas et al., 2017; Simonetti et al., 2019) or indirectly via cargo adaptors (Strochlic et al., 2007; Lauffer et al., 2010; Harterink et al., 2011; Temkin et al., 2011; Steinberg et al., 2012; Steinberg et al., 2013; Gallon et al., 2014; McNally et al., 2017). Working alongside these complexes, the endosome-associated Wiscott-Aldrich syndrome protein and SCAR Homologue (WASH) complex drives the ARP2/3-mediated formation of branched F-actin networks (Derivery et al., 2009; Gomez and Billadeau, 2009). Together, cargo recognition and organisation of a localised F-actin network leads to the formation of one or more retrieval sub-domains on the cytosolic face of the endosomal membrane that provide platforms for the co-ordinated biogenesis of cargo-enriched transport carriers (Puthenveedu et al., 2010).

In higher metazoans, retromer is a stable heterotrimer of VPS35, VPS29 and VPS26 (mammals express two paralogs VPS26A or VPS26B) (Burd and Cullen, 2014). Retromer is associated to endosomes through binding to sorting nexin-3 (SNX3) (Harterink et al., 2011), RAB7-GTP (Rojas et al., 2008; Seaman et al., 2009) and by association to cargo presented sorting motifs (Harrison et al., 2014; Lucas et al., 2016). Retromer also binds to sorting nexin-27 (SNX27), a cargo adaptor for the sequence-dependent inclusion of around 400 cargo proteins that contain a carboxy-terminal PDZ-binding motif (Temkin et al., 2011; Steinberg et al., 2013; Gallon et al., 2014; Clairfeuille et al., 2016). Retromer’s principal role therefore is to orchestrate the retrieval of hundreds of internalised cargo and to promote their recycling to the cell surface (Chen et al., 2010; Temkin et al., 2011; Steinberg et al., 2013). That said, controversy remains as to the role of retromer in a distinct retrieval pathway, the retrograde endosome-to-TGN sorting of the cation-independent mannose 6-phosphate receptor (CI-MPR) (reviewed in Seaman, 2018).

At steady state, CI-MPR is enriched at the TGN where it associates with newly synthesised hydrolase precursors (Braulke and Bonifacino, 2009). The resulting CI-MPR:hydrolase complex is transported to the endosomal pathway, where the acidified endosomal lumen induces the release of the hydrolase. While the hydrolase precursors are delivered to the lysosome, where they contribute to the degradative capacity of this organelle, the unoccupied CI-MPR is retrieved and recycled to the TGN for further rounds of hydrolase delivery. Many studies in mammalian cells are consistent with retromer in regulating CI-MPR transport (Arighi et al., 2004; Seaman, 2004; Seaman, 2007; Wassmer et al., 2007; Bulankina et al., 2009; Harbour et al., 2010; Hao et al., 2013; Breusegem and Seaman, 2014; McGough et al., 2014; Cui et al., 2019). However, we and others have recently questioned this model (Kvainickas et al., 2017; Simonetti et al., 2017). Rather, structural, biochemical and functional evidence has associated ESCPE-1 in sequence-dependent endosome-to-TGN CI-MPR transport through direct recognition of a bipartite sorting motif (Kvainickas et al., 2017; Simonetti et al., 2017; Simonetti et al., 2019).

Part of this controversy may stem from technical variability and in particular the reliance on the generation of retromer knockdown and knockout cells (Seaman, 2018). These procedures induce the gradual loss of retromer expression over the course of hours and days, a time window that has the potential to initiate the activation of compensatory pathways that suppress phenotypes or result in variable and subtle phenotypes. Here we have applied the ‘knocksideways’ methodology (Robinson et al., 2010) to acutely remove retromer and trap this complex on an organelle not implicated in retromer function. Using time-resolved analysis of cargo trafficking, we show that while acute retromer inactivation leads to robust defects in the endosomal recycling of the prototypical retromer cargo GLUT1, no detectable perturbation was observed in the distribution of the CI-MPR. In contrast, acute knocksideways mediated inactivate of ESCPE-1 results in perturbed endosome-to-TGN transport of the CI-MPR. Our study therefore defines a method for the acute inactivation of endosomal retrieval and recycling complexes and provides further data to support the need to reflect on the central role of retromer in the retrograde sorting from the endosome to the *trans*-Golgi network.

## RESULTS

### Retromer knocksideways design and temporal dynamics

To design the retromer knocksideways, we first engineered a cassette encoding for the core VPS35 subunit fused through a carboxy-terminal flexible linker to green fluorescent protein (GFP), which was itself linked to the amino-terminus of rapamycin-binding (FRB) domain (resultant construct encode VPS35-GFP-FRB, Figure 1A). In light of evidence linking retromer to aspects of lysosomal function (*e.g.* Kvainickas et al., 2019), we utilised a modified version of FRB (T2098L) that enables the induction of heterodimerisation by rapalog (AP21967), a compound that has a lower affinity to endogenous mTOR than rapamycin (Clackson et al., 1998). To validate that the VPS35-GFP-FRB chimera was functional we expressed the construct in a previously characterised VPS35 knockout HeLa cell line (Kvainickas et al., 2017). The VPS35-GFP-FRB chimera localised to cytosolic puncta that were identified as endosomes by means of colocalization with the endosome marker EEA1 (Supplementary Figure 1A). GFP-nanotrap immunoprecipitation confirmed that VPS35-GFP-FRB retained the ability to associate with the core retromer components VPS26 and VPS29 (Supplementary Figure 1B). Consistent with retromer assembly, expression of the VPS35-GFP-FRB chimera in the VPS35 knockout HeLa cells reverted the observed lysosomal missorting of GLUT1 and allowed recycling of the transporter back to the cell surface (Supplementary Figure 1C and D). The designed VPS35-GFP-FRB chimera is therefore correctly localised to endosomes and displays the ability to assemble into a retromer complex that retains its function in endosomal cargo retrieval and recycling.

**Figure 1.**
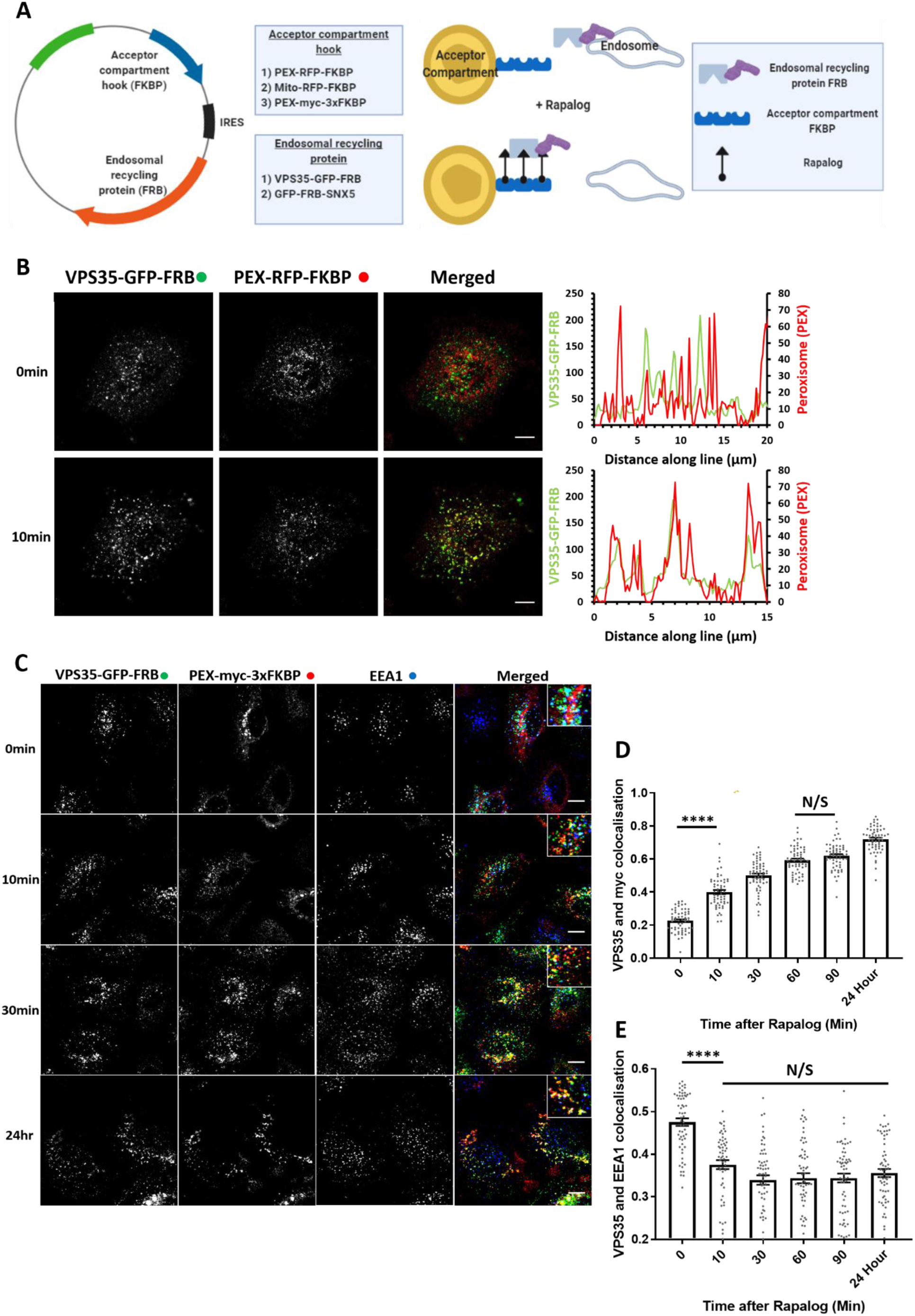
Knocksideways can rapidly mislocalise retromer from endosomes. (A) Schematic showing the design of the endosomal knocksideways system. (B) HeLa cells transfected with retromer knocksideways (PEX-RFP-FKBP and VPS35-GFP-FRB). Still frames are shown from a movie (Supplementary Movies 1A) at either 0 minutes or 10 minutes after the addition of rapalog. Line scans were generated using ImageJ drawing a line through peroxisome structures and represent the co-localization between VPS35-GFP-FRB and PEX-RFP-FKBP at each timepoint. The merged panel displays both channels. Scale bars = 10 µm. (C) Retromer knocksideways HeLa cells were fixed at multiple time points after the addition of rapalog. Anti-myc and anti-EEA1 were used to label PEX-myc-3xFKBP and early endosomes respectively and the merged panel displays triple colocalization between three channels. Zoom panels are displayed in the top right of the merged image. Scale bars = 10 µm. (D) Pearsons co-localization between VPS35-GFP-FRB and PEX-myc-3xFKBP (peroxisomal targeting sequence) at multiple timepoints after the addition of rapalog. *n*_exp_ = 3, *n*_cell_ = 60 with all data points being displayed. Statistical analysis performed - Ordinary one-way ANOVA with multiple comparisons, ****<0.0001 and N/S >0.05. (E) Pearsons co-localization between VPS35-GFP-FRB and EEA1 (early endosome marker) at multiple time points after the addition of rapalog. *n*_exp_ =3, n=60 with all data points being displayed. Statistical analyse performed - Ordinary one-way ANOVA with multiple comparisons, ****<0.0001 and N/S >0.05.

To engineer the acceptor compartment, we fused red fluorescent protein (RFP) to FKBP and linked this to either a mitochondrial targeting sequence (yeast Tom70p – forming Mito-RFP-FKBP (Robinson et al., 2010) or a peroxisomal targeting sequence (PEX3 (residues 1-42) – forming PEX-RFP-FKBP (Kapitein et al., 2010)). To ensure a balanced co-expression, we cloned the genes encoding for Mito-RFP-FKBP and VPS35-GFP-FRB into a bicistronic vector and generated a corresponding bicistronic vector for PEX-RFP-FKBP and VSP35-GFP-FRB (Figure 1A). To visualise the temporal dynamics of retromer knocksideways we performed live imaging immediately after the application of rapalog. For both the mitochondrial and peroxisomal knocksideways systems we observed dynamic accumulation of VPS35-GFP-FRB onto the corresponding acceptor compartment (Supplementary Movies 1A and B), such that approximately 10 minutes after induction of dimerization there was clear co-localization between retromer and the acceptor compartment (Figure 1B and Supplementary Figure 1E).

Considering that retromer has been implicated in mitochondrial function (Braschi et al., 2010; Tang et al., 2015; Wang et al., 2016), we decided to focus on developing the peroxisomal acceptor compartment system: to date, peroxisomes have not been implicated in retromer biology. To increase the capacity of the acceptor compartment we converted PEX-RFP-FKBP to PEX-Myc-3xFKBP (each FKBP separated by flexible linkers GGSGGGSGGAP) (Figure 1A). In transiently transfected HeLa cells, the PEX-Myc-3xFKBP chimera displayed colocalization with the known peroxisome marker PMP70 (Supplementary Figure 1F).

In VPS35 knockout HeLa cells transiently transfected to express PEX3-myc-3xFKBP and VPS35-GFP-FRB, the addition of 100 nM of rapalog established that rerouting of VPS35-GFP-FRB from EEA1-positive endosomes to peroxisomes was achieved within 10 minutes and was complete by 30 minutes (Figure 1C-E) - in the continued presence of rapalog the peroxisome rerouted VPS35-GFP-FRB was retained on this organelle (maximum time studied 24 hours). Together these data establish a method for the acute knocksideways of a functional VPS35-GFP-FRB construct.

### Using knocksideways to examine *in cellulo* retromer assembly

GFP-nanotrap immunoisolation is an established method for identifying protein-protein interactions, including those of the retromer complex (McGough et al., 2014; McMillan et al., 2016). Here we used knocksideways to analyse protein-protein interactions *in cellulo.* Consistent with the assembly of VPS35-GFP-FRB into a functional heterotrimeric complex (Supplementary Figure 1B), analysis of the endogenous localisation of VPS26 revealed that it too was rerouted to peroxisomes with a similar kinetic profile to that observed for VPS35-GFP-FRB (the lack of a suitable antibody precluded the equivalent analysis of VPS29) (Figure 2A and B). In addition, the major retromer accessory complex, the FAM21 containing WASH complex (Derivery et al., 2009; Gomez and Billadeau, 2009; Harbour et al., 2012; Jia et al., 2012) was also rerouted to peroxisomes upon retromer knocksideways (Figure 2C and D, Supplementary Figure 2A and B). Supporting evidence that a sub-population of the WASH complex is associated with endosomes independent of retromer (McNally et al., 2017; MacDonald et al., 2018), a significant amount of the WASH complex was retained on endosomes even after retromer knocksideways (Figure 2C and E, Supplementary Figure 2A, C and D). Retromer knocksideways was selective in that VPS35L, the core component of the functionally distinct retriever complex (McNally et al., 2017), retained its endosome association and was not recruited to peroxisomes upon retromer knocksideways (Supplementary Figure 2E-G).

**Figure 2.**
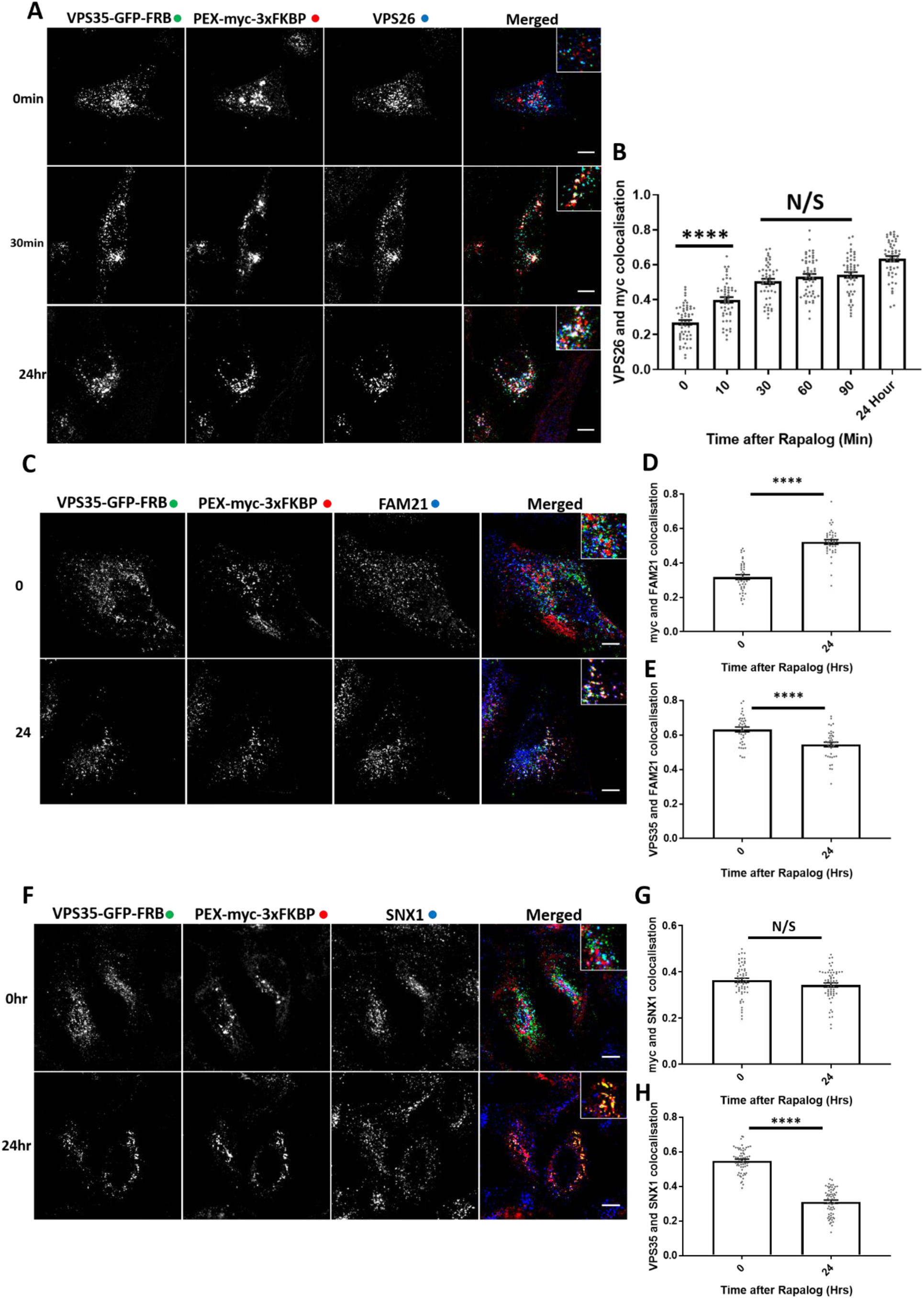
Retromer knocksideways ‘drags’ biochemically validated interacting proteins onto peroxisomes. (A) Retromer knocksideways HeLa cells were fixed before or after multiple time points of rapalog addition and labelled for anti-myc and anti-VPS26. A merged panel displays all three channels combined with a zoom panel. Scale bars = 10 µm. (B) Pearsons colocalization between VPS26 and myc at multiple timepoints, *n*_exp_ = 3, *n*_cell_ = 49-54 with all datapoints being displayed. Statistical analysis performed - Ordinary one-way ANOVA with multiple comparisons, ****<0.0001 and N/S >0.05. (C) Retromer knocksideways HeLa cells were fixed before or after 24 hours of rapalog addition and then labelled with anti-myc and anti-FAM21. The merged panel displays all three channels with zoom-in panels. Scale bars = 10 µm. (D) Pearsons colocalization between anti-myc and anti-FAM21 before and after 24 hours of rapalog, *n*_exp_ = 2, *n*_cell_ = 40 with all data points being displayed. Statistical analysis performed Welch’s t-test ****<0.0001. (E) Pearsons colocalization between anti-VPS35 and anti-FAM21 before and after 24 hours of rapalog, *n*_exp_ = 2, *n*_cell_ = 40 with all data points being displayed. Statistical analysis performed – Welch’s t-test ****<0.0001. (F) Retromer knocksideways HeLa cells were fixed before or after 24 hours of rapalog and then labelled for anti-myc and anti-SNX1. The merged panel displays the PEX-myc-3xFKBP and SNX1 channels combined and has a zoom panel. Scale bars = 10 µm. (G) Pearsons co-localization between myc and SNX1 before and after the addition of rapalog for 24 hours, *n*_exp_ = 3, *n*_cell_ = 60 with all datapoints being displayed. Statistical analysis performed - Welch’s t-test ****<0.0001 (H) Pearsons co-localization between VPS35-GFP-FRB and SNX1 before and after the addition of rapalog for 24 hours, *n*_exp_ = 3, *n*_cell_ = 60 with all datapoints being displayed. Statistical analysis performed - Welch’s t-test N/S>0.05.

Given the selectivity of retromer knocksideways we also decided to apply this methodology to examine the *in cellulo* relationship between retromer and the SNX-BAR proteins that assemble to form the ESCPE-1 complex (Simonetti et al., 2019). In yeast these SNX-BAR proteins associate with the Vps26:Vps35:Vps29 heterotrimer to form the stable pentameric retromer complex (Seaman et al., 1998). In metazoans however, retromer and ESCPE-1 appear to function independently, which is inconsistent with the formation of a long-lived and stable pentameric complex (Kvainickas et al., 2017; Simonetti et al., 2017). Indeed, we failed to observe the rerouting of endogenous SNX1, a component of the ESCPE-1 complex, onto peroxisomes after 24 hours of rapalog treatment (Figure 2F-H). These data therefore support the *in cellulo* and *in vivo* evidence that in metazoans retromer and ESCPE-1 have evolved into functionally distinct complexes (Kvainickas et al., 2017; Simonetti et al., 2017; Simonetti et al., 2019; Strutt et al., 2019). Overall, the designed VPS35 knocksideways provides a method for the acute and selective rerouting of retromer (and its functional coupled accessory proteins) away from endosomes to the neighbouring peroxisome organelle.

### Acute retromer knocksideways leads to a time-resolved GLUT1 sorting defect

Retromer and retromer-associated cargo adaptors have been shown to control the endosomal retrieval and recycling of >400 cell surface proteins including the glucose transporter GLUT1 (Steinberg et al., 2013). To define the functional consequences of retromer knocksideways we examined the steady-state distribution of GLUT1 in VPS35 knockout HeLa cells rescued by expression of VPS35-GFP-FRB knocksideways. Following the addition of rapalog for 24 hours, fixed cell confocal imaging revealed a GLUT1 missorting phenotype, defined by the steady-state loss of GLUT1 at the cell surface and the enrichment of GLUT1 with LAMP1-positive late endosomes / lysosomes (Figure 3A and B). To time-resolve the appearance of the GLUT1 trafficking phenotype, we fixed cells at various time points following rapalog addition. Quantification established that a statistically significant GLUT1 missorting phenotype began to emerge after 1-3 hours of retromer knocksideways and reached a maximum penetrance after 10 hours (Figure 3C and D). The difference between the time scales of retromer knocksideways (Figure 1C-E) compared with the appearance of the GLUT1 missorting phenotype is entirely consistent with the known rate of GLUT1 lysosomal mediated degradation observed upon retromer suppression and reflects the relatively slow rate of endocytosis of this transporter (Steinberg et al., 2013).

**Figure 3.**
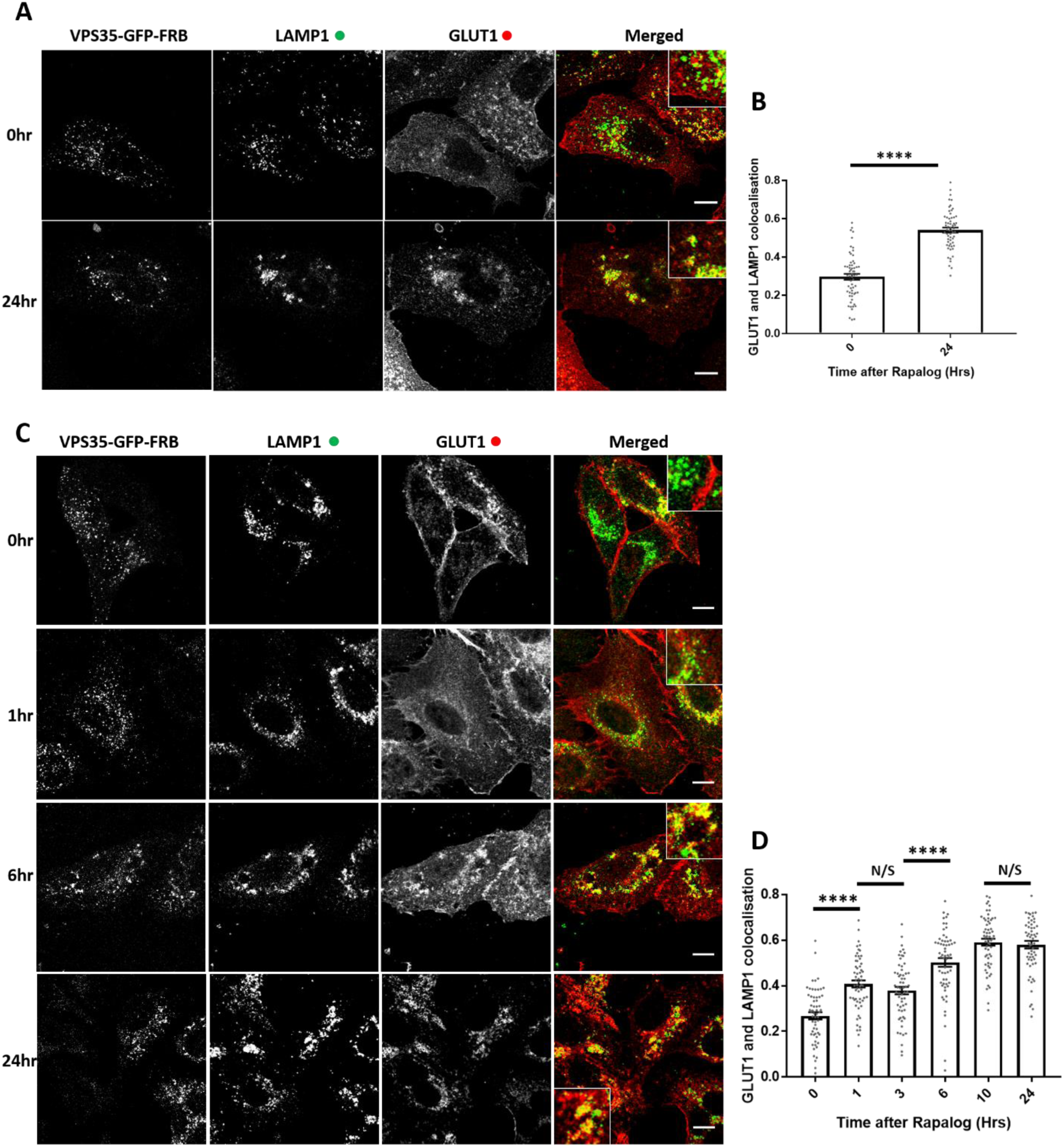
Retromer knocksideways results in the rapid functional inactivation of retromer and the temporal resolution of the accumulation of retromer depleted phenotypes. (A) Retromer knocksideways HeLa cells were fixed before or after 24 hours of rapalog addition. Anti-LAMP1 and anti-GLUT1 were then used to label the late endosome/lysosome and retromer cargo respectively. The merge panel displays both the LAMP1 (green) and the GLUT1 (red) channels with a zoom panel. Scale bars = 10 µm. (B) Pearsons colocalization between GLUT1 and LAMP1 before and after 24 hours, *n*_exp_ = 3, *n*_cell_ = 60 with all data points being displayed. Statistical analysis performed - Welch’s t-test ****<0.0001. (C) Retromer knocksideways HeLa cells were fixed before and after multiple time points of rapalog addition. Anti-LAMP1 and anti-GLUT1 were then used to label late endosome/lysosome and retromer cargo respectively. The merge panels display both the LAMP1 and GLUT1 labelling with a zoom panel. Scale bars = 10 µm. (D) Pearsons colocalization between GLUT1 and LAMP1 before and after multiple timepoints of rapalog, *n*_exp_ = 3, *n*_cell_ = 60 with all data points being displayed. Statistical analyses performed - Ordinary one-way ANOVA with multiple comparisons, ****<0.0001 and N/S >0.05.

The missorting of GLUT1 upon retromer knocksideways was not the result of a global effect on endosomal sorting as the endosomal retrieval and recycling of the retriever-dependent cargo, α_5_β_1_-integrin was not affected upon retromer knocksideways (Supplementary Figure 3A and B) - consistent with the lack of effect of retromer knocksideways on the endosomal association of the retriever complex (Supplementary Figure 2E-G). Moreover, the development of the GLUT1 missorting did not stem from the recruiting of ‘foreign’ proteins to peroxisomes as retromer knocksideways performed in wild-type HeLa cells, which retain expression of endogenous VPS35 that is not subject to knocksideways, did not illicit the development of a GLUT1 missorting phenotype (Supplementary Figure 3C and D). Together, these data support that it is the specific removal and inactivation of retromer that causes the time-resolved development of the observed GLUT1 missorting phenotype.

### Retromer-independent CI-MPR retrograde trafficking

Next, we investigated the role of retromer in the retrograde trafficking of CI-MPR from endosomes to the TGN (Arighi et al., 2004; Seaman, 2004; Kvainickas et al., 2017; Simonetti et al., 2017; Cui et al., 2019). In VPS35 knockout HeLa cells rescued through expression of VPS35-GFP-FRB, the CI-MPR is chiefly localized to the perinuclear TGN as defined through co-localization with TGN markers Golgin97 and TGN46 (Figure 4A and B, Supplementary Figure 4A and B). After the addition of rapalog and initiation of retromer knocksideways we failed to observe a detectable alteration in the steady-state distribution of the CI-MPR (Figure 4A and B, Supplementary Figure 4A and B) over a time frame where the endosomal missorting of internalized GLUT1 was readily observed (Figure 3C and D). Given that the endosome-to-TGN transport of the CI-MPR is considered to occur over a period of approximately 20 - 30 minutes (Seaman, 2004), the acute perturbation of retromer function does not appear to lead to a detectable defect in the endosomal sorting of the CI-MPR. These data are consistent with a limited role for retromer in endosome-to-TGN retrograde transport of this receptor (Kvainickas et al., 2017; Simonetti et al., 2017).

**Figure 4.**
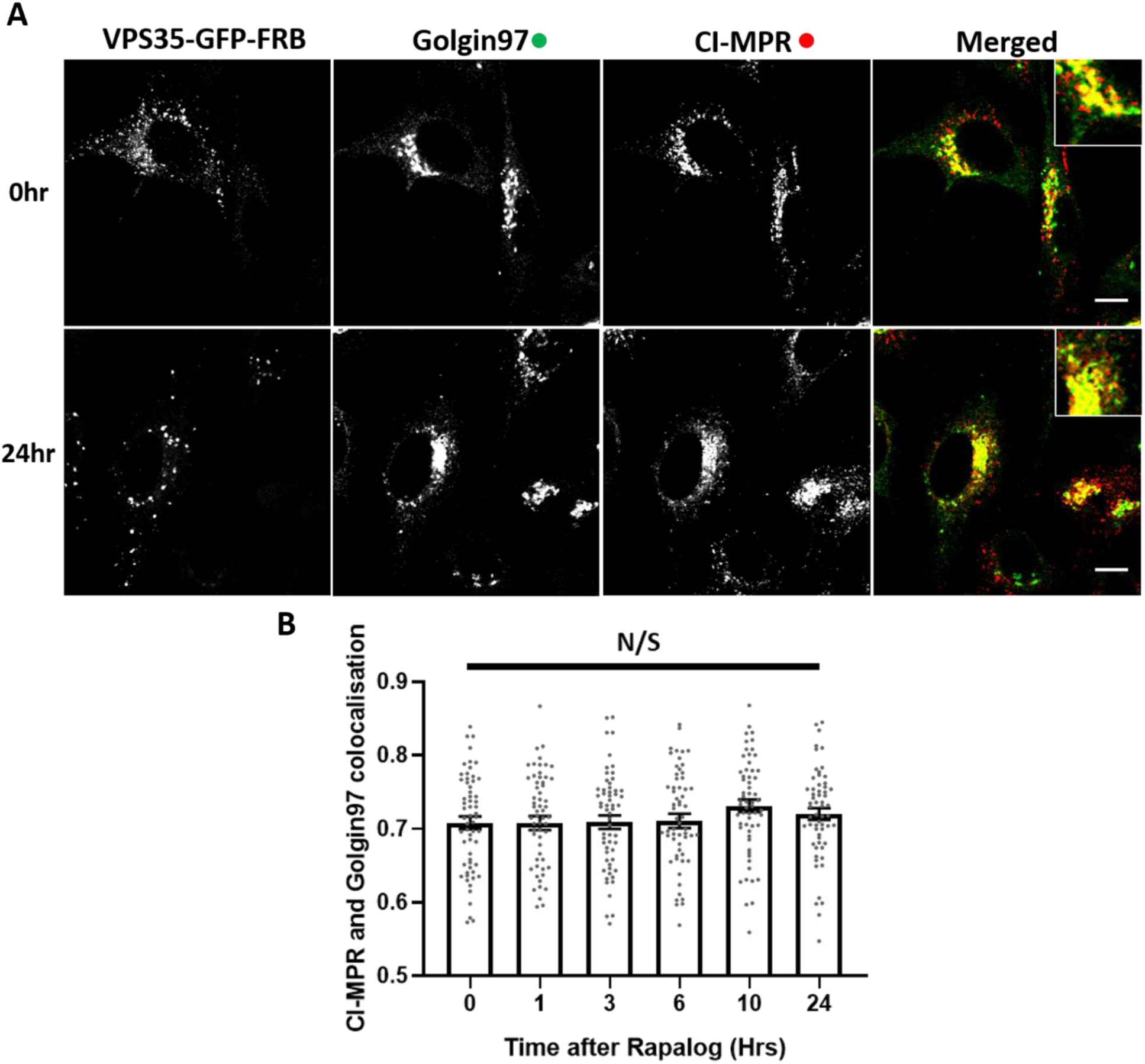
Knocksideways indicates no visible role for retromer in SNX-BAR mediated retrograde transport of CI-MPR. (A) Retromer knocksideways HeLa cells were fixed before or after multiple timepoints of rapalog addition and labelled for Golgin97 and CI-MPR. A merged panel displays both the anti-Golgin97 and anti-CI-MPR channels and also has a zoom panel. Scale bars = 10 µm. (B) Pearsons colocalization between CI-MPR and Golgin97 at multiple timepoints, *n*_exp_ = 3, *n*_cell_ = 60 with all datapoints being displayed. Statistical analysis performed - Ordinary one-way ANOVA with multiple comparisons, N/S >0.05.

### Knocksideways of ESCPE-1 leads to a time-resolved defect in CI-MPR sorting

The ESCPE-1 complex regulates sequence-dependent endosome-to-TGN transport of the CI-MPR (Simonetti et al., 2019). ESCPE-1 comprises a heterodimer of SNX1 or SNX2 (these proteins are functionally redundant) associated with either SNX5 or SNX6, which are also functionally redundant (Wassmer et al., 2009). Of these proteins, it is the PX domains of SNX5 and SNX6 that directly bind to the ΦxΩxΦ(x)nΦ sorting motif in CI-MPR to mediate endosome-to-TGN transport (Simonetti et al., 2019). To provide a positive control for the lack of detectable effect of retromer knocksideways on CI-MPR trafficking we therefore constructed a bicistronic vector encoding for PEX3-myc-3xFKBP and GFP-FRB-SNX5 (Figure 1A). When expressed in HeLa cells SNX5-GFP-FRB localised to endosomes as defined by co-localization with EEA1 (Supplementary Figure 5A). Interestingly, after rapalog addition we observed a slight recruitment of EEA1 to the peroxisomal hook indicating a movement of the endosomal compartment to the peroxisomal compartment (Figure 5A and B). However, this ‘dragging’ of endosomes was not complete as there was still a loss of co-localization between GFP-FRB-SNX5 and EEA1 (Figure 5A and C). In GFP-FRB-SNX5 knocksideways endogenous SNX1 was recruited to peroxisomes after rapalog treatment with no loss of co-localization between SNX5 and SNX1 indicating a recruitment of the functional ESCPE-1 complex (Figure 5D-F).

**Figure 5.**
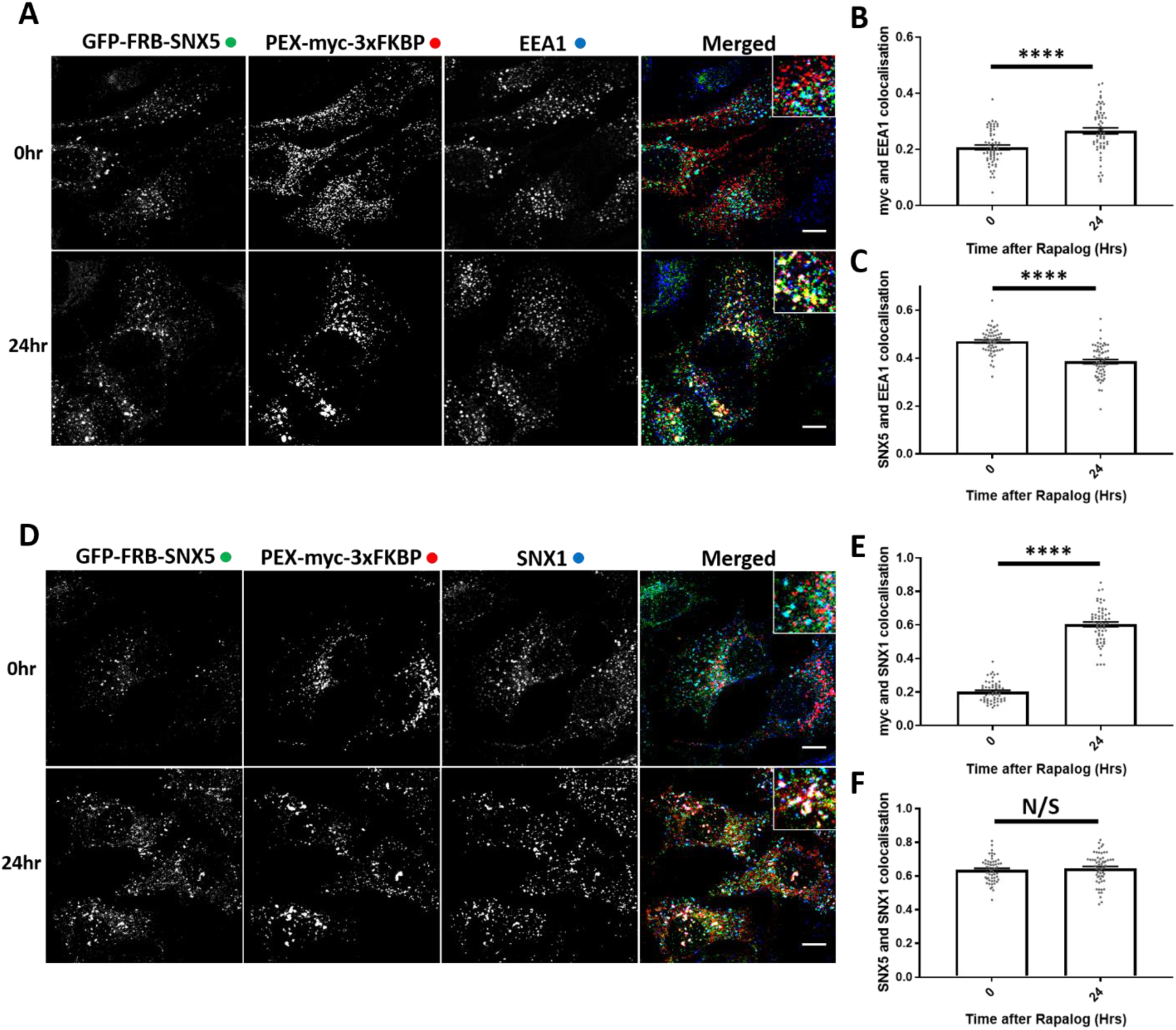
ESCPE-1 knocksideways results in the dragging of a small population of endosomes and peroxisomes together whilst still removing ESCPE-1 from endosomes. (A) ESCPE-1 knocksideways HeLa cells were fixed before and after the addition of rapalog for 24 hours and then labelled for anti-myc and anti-EEA1. The merged panel shows all three channels with a zoom panel. Scale bars = 10 µm. (B) Pearsons co-localization between myc and EEA1 before and after 24 hours of rapalog, *n*_exp_ = 3, *n*_cell_ = 60 with all data points being displayed. Statistical analysis performed - Welch’s t-test, ****<0.0001 (C) Pearsons co-localization between GFP-FRB-SNX5 and EEA1 before and after 24 hours of rapalog, *n*_exp_ = 3, *n*_cell_ = 60 with all data points being displayed. Statistical analysis performed – Welch’s t-test, ****<0.0001. (D) ESCPE-1 knocksideways HeLa cells were fixed before and after the addition of rapalog for 24 hours and then labelled for anti-myc and anti-SNX1. The merged panel shows all three channels with a zoom panel. Scale bars = 10 µm. (E) Pearsons co-localization between myc and SNX1 before and after 24 hours of rapalog, *n*_exp_ = 3, *n*_cell_ = 53-60 with all data points being displayed. Statistical analysis performed – Welch’s t-test, ****<0.0001. (F) Pearsons co-localization between GFP-FRB-SNX5 and SNX1 before and after 24 hours of rapalog, *n*_exp_ = 3, *n*_cell_ = 53-60 with all data points being displayed. Statistical analysis performed – Welch’s t-test, N/S >0.05.

Next we used the GFP-FRB-SNX5 knocksideways system to time-resolve CI-MPR endosome-to-TGN trafficking. Expression of the GFP-FRB-SNX5 chimera in a previously isolated and characterised SNX5/SNX6 double knockout HeLa cell line (Simonetti et al., 2017) reverted the observed missorting of the CI-MPR and allowed the receptor to return to its normal steady-state localisation (Supplementary Figure 6A and B). Consistent with the role of SNX5 in the ESCPE-1-mediated endosome-to-TGN transport of the CI-MPR (Kvainickas et al., 2017; Simonetti et al., 2017; Simonetti et al., 2019), SNX5 knocksideways in SNX5/SNX6 double knockout HeLa cells led to the time-resolved appearance of a CI-MPR missorting phenotype as defined by a reduced enrichment of the CI-MPR at the Golgin97 or TGN46-labelled TGN with a maximum penetrance at 6 hours (Figure 6A and B, Supplementary Figure 6C and D). There was no observed defect in α_5_β_1_-integrin recycling during GFP-FRB-SNX5 knocksideways indicating the selective nature of this procedure (Figure 6C and D). Together these data establish that acute perturbation of the ESCPE-1 complex leads to a missorting of CI-MPR, thereby reinforcing the functional significance of the lack of observed effect of retromer knocksideways on the trafficking of this receptor.

**Figure 6.**
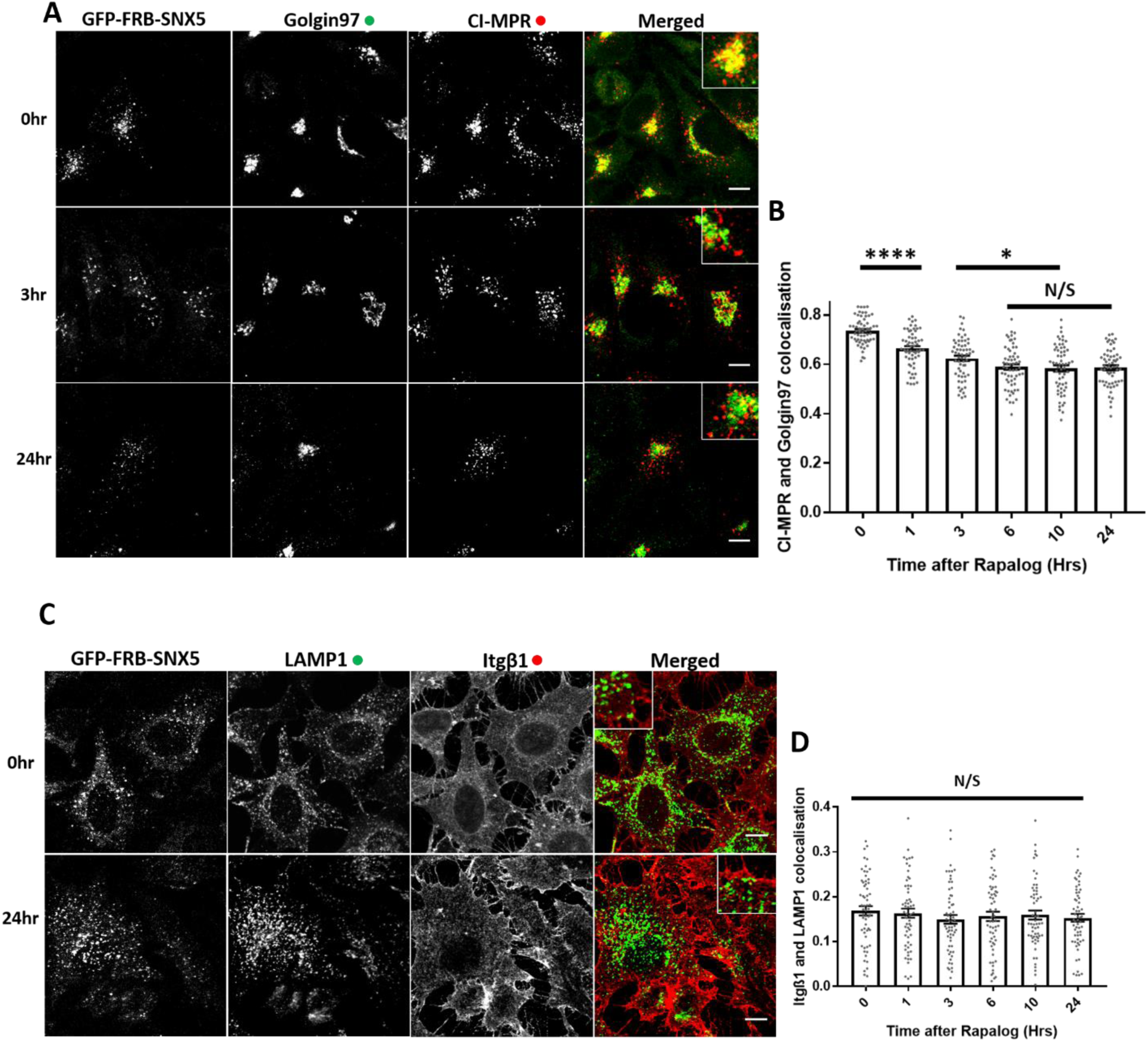
ESCPE-1 knocksideways inactivates ESCPE-1 and results in a temporally resolved CI-MPR redistribution away from the TGN. (A) SNX-BAR knocksideways HeLa cells were fixed before and after the addition of rapalog for multiple timepoints and then labelled for anti-Golgin97 and anti-CI-MPR. The merged panel shows both the anti-Golgin97 and anti-CI-MPR channels with a zoom panel. Scale bars = 10 µm. (B) Pearsons co-localization between Golgin97 and CI-MPR before and after multiple timepoints of rapalog, *n*_exp_ = 3, *n*_cell_ = 60 with all data points being displayed. Statistical analysis performed - Ordinary one-way ANOVA with multiple comparisons, ****<0.0001, *<0.05, N/S >0.05. (C) SNX-BAR knocksideways HeLa cells were fixed before and after the addition of rapalog for multiple timepoints and then labelled for anti-LAMP1 and anti-Itgβ1. The merged panel shows both the LAMP1 and Itgβ1 channels with a zoom panel. Scale bars = 10 µm. (D) Pearsons co-localization between LAMP1 and Itgβ1 before and after multiple timepoints of rapalog, *n*_exp_ = 3, *n*_cell_ = 60 with all data points being displayed. Statistical analysis performed - Ordinary one-way ANOVA with multiple comparisons, N/S >0.05.

### H4 human neuroglioma knocksideways is consistent with a limited role for retromer in retrograde CI-MPR trafficking

Our study of endosomal cargo sorting associated with depletion or knocksideways of sorting machinery has so far been limited to a single non-neuronal cell type. To extend these observations, we therefore turned to H4 neuroglioma cells and generated both retromer knockout (targeting VPS35) and ESCPE-1 knockout cells (dual targeting of SNX5 and SNX6). Interestingly, confocal imaging of the retromer knockout cells revealed an enhanced intensity in the staining of endogenous CI-MPR that was not observed in the ESCPE-1 knockout cells (Figure 7A and B). Despite the increase in the CI-MPR signal intensity, retromer knockout cells did not display a significant change in the quantified Pearsons correlation coefficient between CI-MPR and Golgin97. This is consistent with a limited role for retromer in CI-MPR retrograde trafficking in H4 neuroglioma cells (Figure 7A and C). In contrast, the ESCPE-1 knockout H4 neuroglioma cells displayed a clear redistribution of CI-MPR to peripheral dispersed puncta (Figure 7A, D and Supplementary Figure 7A).

**Figure 7.**
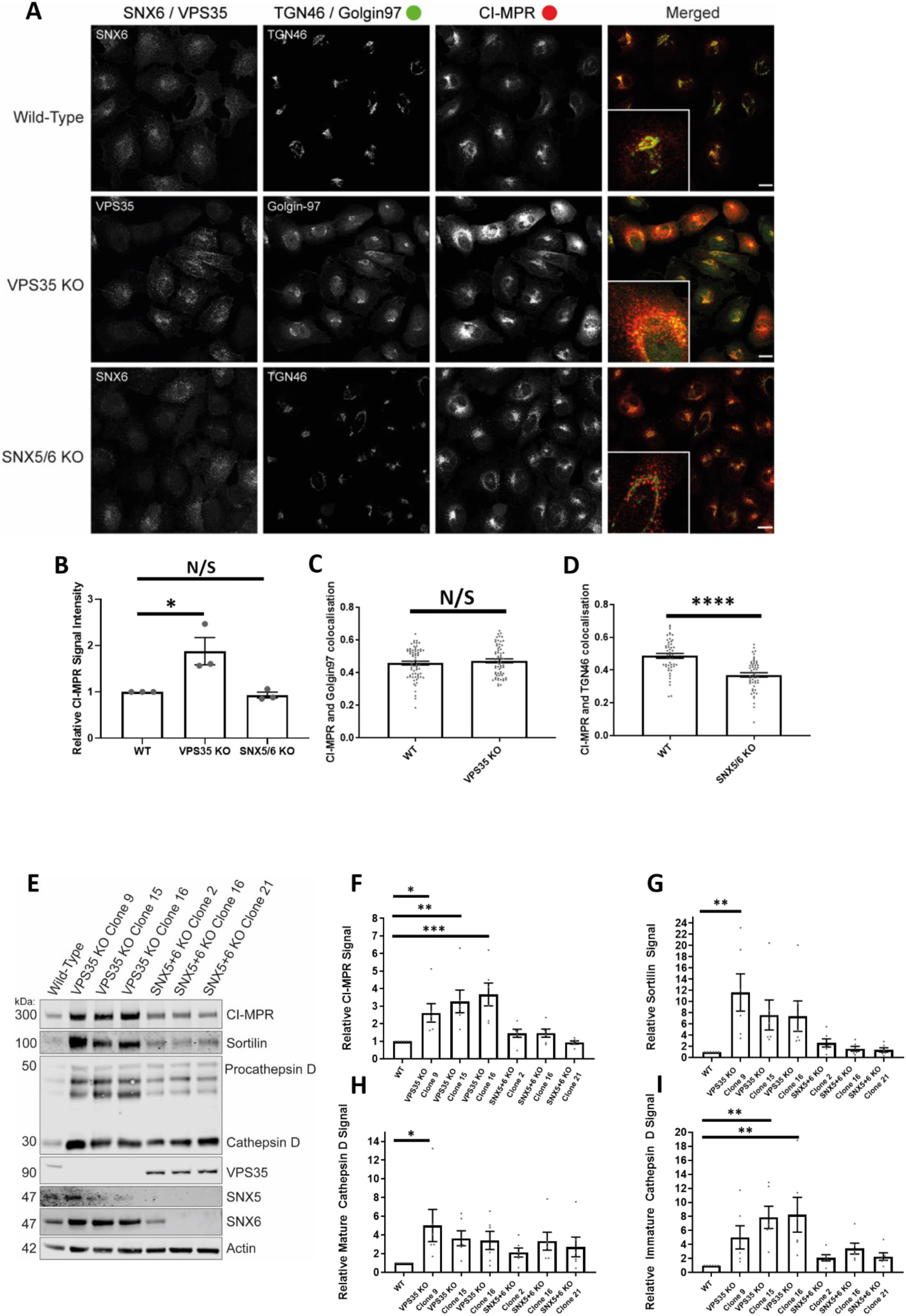
VPS35 knockout H4 neuroglioma cells display an upregulation of lysosomal hydrolases and lysosomal hydrolase receptors. (A) VPS35 and SNX5/SNX6 dual knockout mixed population H4 neuroglioma cells were generated and then fixed. Cells were stained with either anti-VPS35 or anti-SNX6 to confirm which cells were knocked out in the mixed population. Cells were also co-stained with both anti-CI-MPR and either anti-TGN46 (SNX5/SNX6 dual knockout) or anti-Golgin97 (VPS35 knockout). The merged panel displays both the CI-MPR and TGN46 or Golgin97 channels. Scale bars = 20 µm. (B) Normalised values for relative CI-MPR signal intensity between conditions, *n*_exp_ = 3, *n*_cell_ = 44-68 with average value data being shown for each M. Statistical analysis performed - Students t-test *<0.05, N/S >0.05. (C) Pearsons co-localization between CI-MPR and Golgin97 in wild-type and VPS35 knockout cells, *n*_exp_ = 3, *n*_cell_ = 64-69 with all data points being displayed. Statistical analysis performed - Welch’s t-test, N/S >0.05. (D) Pearsons co-localization between CI-MPR and TGN46 in wild-type and SNX5/SNX6 knockout cells, *n*_exp_ = 3, *n*_cell_ = 49-53 with all data points being displayed. Statistical analysis performed - Welch’s t-test, ****<0.0001. (E) Western blot analysis of wild-type and VPS35 knockout or SNX5/SNX6 knockout clonal cell lines using anti-CI-MPR, anti-Sortilin, anti-Cathepsin D, anti-VPS35, anti-SNX5, anti-SNX6, anti-Actin. (F-I) Relative (actin) measured signals for wild-type and VPS35 knockout or SNX5/SNX6 dual knockout clonal cell lines for CI-MPR, Sortilin, mature Cathepsin D, immature Cathepsin D. Statistical analysis performed - Ordinary one-way ANOVA with multiple comparisons, ***<0.001, **<0.01, *<0.05.

To extend these data, we isolated individual clonal retromer and ESCPE-1 knockout H4 cell lines. Biochemical analysis of 3 independent clonal lines revealed that retromer knockout resulted in a pronounced up-regulation of CI-MPR expression (Supplementary Figure 7B). Moreover, the expression of another lysosomal hydrolase receptor, sortilin, was also increased across all three independent lines as was the immature and mature forms of the lysosomal hydrolase cathepsin D (Figure 7E-I). These increases in protein expression were not observed across 3 independent ESCPE-1 knockout H4 cell lines (Figure 7E-I).

To examine whether the increased expression of CI-MPR, sortilin and cathepsin D arose from a retromer-dependent trafficking defect, which would be inconsistent with the lack of effect of retromer knockout on the steady state distribution of CI-MPR (Figure 7A and C), or reflected a longer-term compensatory mechanism we established the acute VPS35 knocksideways methodology in H4 cells. Expression of VPS35-GFP-FRB rescued the GLUT1 missorting phenotype in retromer knockout cells. Initiation of VPS35 knocksideways resulted in a time-resolved missorting and accumulation of GLUT1 to LAMP1 positive late endosomes and lysosomes (Figure 8A and B) confirming an acute perturbation in retromer function (Figure 3C and D). In a parallel time-resolved retromer knocksideways we failed to detect a significant redistribution of CI-MPR away from TGN markers Golgin97 and TGN46, consistent with retromer playing a limited role in retrograde trafficking of CI-MPR in H4 cells (Figure 8C and D and Supplementary Figure 8A and B). Together these data are more suggestive of the observed up-regulation in expression of CI-MPR in retromer knockout cells arising as an indirect affect of a compensatory pathway(s) rather than a direct endosome-to-TGN sorting phenotype.

**Figure 8.**
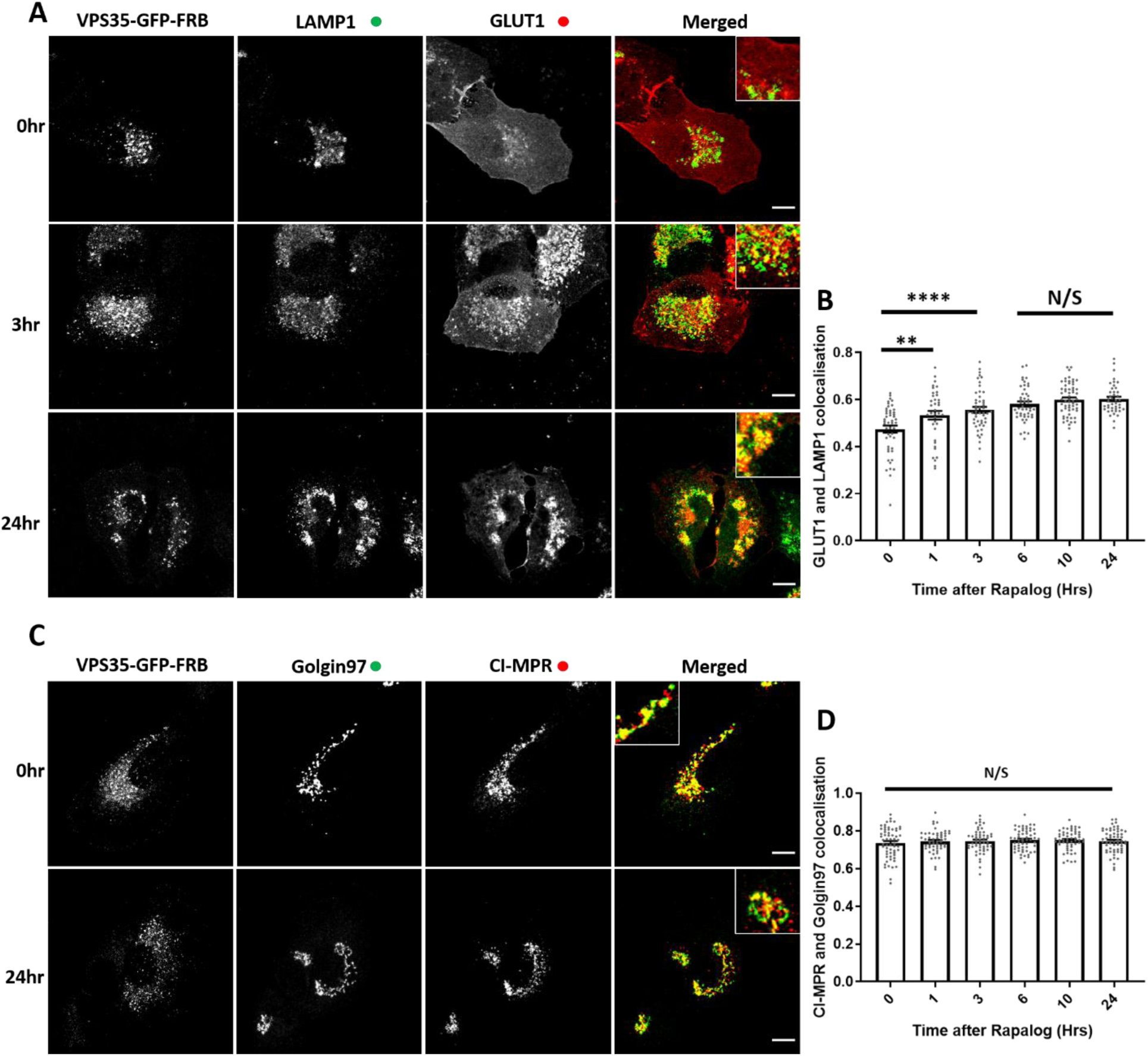
VPS35 knocksideways in H4 neuroglioma cells confirms a role for retromer in recycling of GLUT1 but no visible role for retromer in the retrograde CI-MPR trafficking. (A) Retromer knocksideways H4 neuroglioma cells were fixed before and after multiple time points of rapalog addition. Antibodies towards anti-LAMP1 and anti-GLUT1 were then used to label late endosome/lysosome and retromer cargo respectively. The merge panels display both the LAMP1 and GLUT1 labelling with a zoom-in panel. Scale bars = 10 µm. (B) Pearsons co-localization between GLUT1 and LAMP1 before and after multiple time points of rapalog, *n*_exp_ = 3, *n*_cell_ = 37-58 with all data points being displayed. Statistical analyses performed - Ordinary one-way ANOVA with multiple comparisons, N/S >0.05, **<0.01, ****<0.0001 (C) Retromer knocksideways H4 neuroglioma cells were fixed before or after multiple timepoints of rapalog addition and labelled with anti-Golgin97 and anti-CI-MPR. A merged panel displays both the Golgin97 and CI-MPR channels and has a zoom panel. Scale bars = 10 µm. (D) Pearsons co-localization between CI-MPR and Golgin97 at multiple timepoints, *n*_exp_ = 3, *n*_cell_ = 52-60 with all datapoints being displayed. Statistical analysis performed - Ordinary one-way ANOVA with multiple comparisons, N/S >0.05.

## DISCUSSION

Here we have developed and applied knocksideways to acutely inactivate retromer and the ESCPE-1 complex. Previously developed to inactivate the AP1 and AP2 clathrin adaptors (Robinson et al., 2010; Hirst et al., 2012), this approach provides a method to acutely perturb sorting complex function in a time frame that better aligns with the dynamic nature of endosomal membrane trafficking. By visualising the sorting of endogenous GLUT1 and CI-MPR, our data provide insight into the temporal dynamics of endosomal cargo sorting and support the established role of retromer in cell surface recycling (Chen et al., 2010; Temkin et al., 2011; Steinberg et al., 2013). In contrast, we have provided independent evidence consistent with a comparatively limited role for retromer in ESCPE-1 dependent CI-MPR retrograde sorting in HeLa and H4 human neuroglioma cells (Kvainickas et al., 2017; Simonetti et al., 2017; Simonetti et al., 2019).

In applying knocksideways we have established that retromer and ESCPE-1 can be specifically and rapidly inactivated leading to the time-resolved development of selective cargo sorting defects through the endosomal pathway. Interestingly, in examining CI-MPR phenotypes in H4 neuroglioma cells we have observed a clear distinction between the effects of long-term retromer knockout and acute retromer knocksideways. Only in retromer knockout cells did we observe an increase in the steady-state expression of not only CI-MPR, but also sortillin and cathepsin D. The mechanistic basis of this compensatory up-regulation is presently unclear but it may, in part, reflect indirect effects on the control of RAB7 activity and perturbed mTORC1 signalling leading to induction of autophagy (Jimenez-Orgaz et al., 2018; Kvainickas et al., 2019; Curnock et al., 2019) and the swollen lysosomal phenotype previously documented in retromer knockout cells (Cui et al., 2019).

In contrast to retromer’s pentameric assembly in yeast (Seaman et al., 1998), an increasing body of biochemical, *in cellulo* and *in vivo* functional data point to the fact that the equivalent metazoan assembly has evolved into two functionally distinct complexes, the VPS26:VPS35:VPS29 retromer and the SNX1/SNX2:SNX5/SNX6 ESCPE-1 complex (Norwood et al., 2011; Swarbrick et al., 2011; Kvainickas et al., 2017; Simonetti et al., 2017; Simonetti et al., 2019; Strutt et al., 2019). In utilising knocksideways as an *in cellulo* ‘interaction’ assay we have provided further supporting evidence of the distinct nature of the retromer and ESCPE-1 complexes. Specifically, acute knocksideways of the core VPS35 retromer component results in the equivalent time-resolved knocksideways of the endogenous population of VPS26 but has no detectable effect on the endosomal localisation of ESCPE-1. This technically distinct approach therefore provides further data to support the diversification of retromer and ESCPE-1 into two functionally distinct sorting complexes.

The development of retromer knocksideways has also added to our understanding of the endosomal association of the actin polymerising WASH complex. Direct binding of FAM21 to VPS35 is a major mechanism for the retromer-dependent association of the WASH complex to endosomes (Gomez and Billadeau, 2009; Harbour et al., 2012; Jia et al., 2012). That said, increasing evidence suggests that a sub-population of the WASH complex is associated to endosomes independent of retromer (McNally et al., 2017; Kvainickas et al., 2017; Simonetti et al., 2017; MacDonald et al., 2018). Consistent with these data, acute knocksideways of retromer induces a redistribution of a major proportion of endogenous WASH but a significant sub-population retains an endosomal association.

In summary, by applying knocksideways we have visualized the acute inactivation of retromer and ESCPE-1 and, through quantification of the temporal development of resulting cargo sorting defects, provided clarification of the role of these complexes in the sorting of CI-MPR and GLUT1. Moving forward, we aim to combine knocksideways and our established unbiased proteomic quantification of the cell surface proteome to time-resolve the functional role of endosomal sorting complexes in global cargo retrieval and recycling.

## MATERIALS AND METHODS

### Antibodies

Antibodies used in this study are as follows: SNX1 (clone 51; 611482; BD Bioscience), GLUT1 (ab40084; Abcam), Golgin97 (clone CDF4; A-21270; Thermo Fisher Scientific), VPS26 (ab23892; Abcam), VPS35 (ab10099; Abcam, IF), VPS35 (ab97545; Abcam, IF), VPS35 (ab157220; Abcam, WB), VPS29 (ab98929; Abcam), FAM21 (gifted from D.D. Billadeau, Mayo Clinic, Rochester, MN), EEA1 (N-19; sc-6415; Santa Cruz Biotechnology) TGN46 (AHP500G; Bio-Rad Laboratories), anti-Myc (gifted from H.Mellor, The University of Bristol), LAMP1 (H4A3 was deposited to the DSHB by August, J.T. / Hildreth, J.E.K.; DSHB Hybridoma Product; H4A3), LAMP1 (ab24170; Abcam). Mouse EEA1 (610457; BD Bioscience), CI-MPR (ab124767; Abcam), Itgβ1 (TS2/16), SNX6 (Clone D-5; 365965; Santa Cruz Biotechnology, PMP70 (PA1-650; Invitrogen), WASH1 (gifted from D.D. Billadeau, Mayo Clinic, Rochester, MN), C16orf62 (PA5-28553; Pierce), Sortilin (ab16640; Abcam), Cathepsin D (21327-1-AP, Proteintech), SNX5 (ab180520; Abcam), B-Actin (A1978; Sigma-Aldrich).

### Plasmids

A pIRESneo3 vector was adapted to generate the bicistronic knocksideways system. First, VPS35-GFP and FRB (PCR from the Kinesin-FRB template gifted from L. Kapetein, Utrecht University, Netherlands) were PCR overlapped together inserting a XhoI site between VPS35-GRP and FRB and then ligated downstream of the IRES component between the SmaI and PacI sites. PEX-RFP-FKBP was amplified from the template (gifted from L. Kapetein, Utrecht University, Netherlands) and ligated into the MCS downstream of the CMV promoter in pIRESNEO3 using EcoRV and NotI. The mitochondrial targeting sequence (gift from Scottie Robinson CIMR, UK.) was inserted in place of the PEX targeting sequence using the EcoRV and AgeI restriction sites. To create GFP-FRB-SNX5 first, GFP-FRB was amplified and inserted between the SmaI and PacI sites to generate a new FseI site upstream of the PacI restriction site. The new FseI and PacI site was used to insert SNX5. The PEX-RFP-FKBP was converted to PEX-myc-3xFKBP by PCR of myc-FKBP and inserted between the AgeI and NotI sites to generate PEX-myc-FKBP. The AscI site (upstream of FKBP in the PEX-RFP-FKBP) was used to sequential insert two FKBP cassettes using a MluI-AscI insertion (MluI compatible with AscI but destroying the AscI site allowing the second insertion). CRISPR Cas9 plasmids were obtained from Addgene (#62988, pSpCas9(BB)-2A-Puro PX459 V2.0)

### Cell culture and DNA transfection

HeLa and H4 neuroglioma cells were cultured in DMEM (Sigma) supplemented with 10% (vol/vol) FCS (Sigma) and grown using standard conditions. Lipofectamine LTX was used in DNA transfections. Per six well dish 2 µg of DNA was mixed with 4 µL of the LTX supplement into 100 µL of Opti-Mem. In another incubation 100 µL of Opti-Mem was mixed with 8 µL of Lipofectamine LTX. After a 5-minute incubation, the two 100 µL Opti-Mem mixes were combined and incubated for a further 20 minutes. The 200 µL mix was then added dropwise onto 60-80% confluent HeLa cells and transfected cells were left for 48 hours for DNA expression. VPS35 knockout HeLa cells and SNX5/SNX6 double knockout was generated as previously described and cultured as stated above for wild-type HeLa (Simonetti et al., 2017).

### Generation of H4 clonal cells

H4 cells were seeded the day prior to transfection, then transiently transfected with CRISPR plasmids encoding the Cas9 enzyme, a puromycin resistance marker and specific gRNA guides against VPS35, SNX5 or SNX6 (Kvainickas et al., 2017; Simonetti et al., 2017) using FuGENE® 6 (Promega). The day after transfection, cells are incubated with 1 μg/mL puromycin for 24 hours to select for knockout cells. Following puromycin selection, H4 cells were seeded into a 96-well plate at a density of 1 cell per well in 200 μL Iscove’s Modified Dulbecco’s Medium (Thermo Fisher). Clones were grown to confluency then expanded and screened for successful knockout deletion by Western blotting.

### GFP trap and Western blot analysis

Cells were lysed in GFP trap buffer (50 mM Tris-HCl, 0.5% NP-40 and Roche protease inhibitor cocktail) and the lysate was added to pre-equilibrated GFP trap beads (ChromoTek). Beads were washed 3 times in the GFP trap buffer and then lysates were diluted in 2x sample buffer and boiled for 10 minutes. Proteins were resolved on a NuPAGE 4-12% gels (Invitrogen) and transferred onto polyvinylidene fluoride membrane (EMD Millipore). Once transferred membranes were blocked in TBS 5% milk and the primary antibody (see antibody section) was diluted in TBS 0.1% (v/v) Tween-20 (TBS-T) 5% (w/v) milk and incubated with the membrane for 1 hour at room temperature or overnight at 4°C. Membranes were washed 3 times in TBS-T. Secondary antibodies (see antibody section) were diluted into TBS-T with 5% milk and 0.01% SDS and incubated with the washed membrane for 1 hour at room temperature. TBS-T was used to wash the membrane (3 times) prior to quantitative imaging using an Odyssey scanning system (LI-COR Biosciences). Analysis was performed on Image Studio lite (LI-COR Biosciences).

### Knocksideways

pIRESNEO3 bicistronic vectors encoding the knocksideways peroxisomal/mitochondrial acceptor compartment and either VPS35-GFP-FRB or GFP-FRB-SNX5 were transfected into cells. Either 24 hours of 0.1% (v/v) ethanol vehicle was added to the 0 timepoint or rapalog (100 nM) was added to the 24-hour timepoint. The following day rapalog was added to the different timepoints and then cells were fixed and stained.

### Immunofluorescent staining

Cells were washed once in PBS before fixation for 8 minutes in 4% PFA (16% PFA stock diluted in PBS). Three washes in PBS were performed and then a 5-minute incubation with PBS 100 mM glycine was used to quench the PFA. After three more PBS washes cells were left in PBS overnight. Cells were incubated with PBS + 3% BSA and 0.1% Triton-X100 for 10 minutes and then with PBS + 3% BSA for a further 10 minutes. Primary antibodies (see antibody section) were diluted in PBS + 3% BSA and incubated for 1 hour. Cells were washed 3 times with PBS with the secondary antibody (see antibody section) being diluted into PBS + 3% BSA and incubated for 1 hour. Cells were washed 3 times with PBS and washed once with distilled water before mounting the coverslips in Fluoromount-G.

### Microscopy and image analysis

For image acquisition a Leica SP5-AOBS confocal laser scanning confocal microscope was used attached to a Leica DM I6000 inverted epifluorescence microscope. The 63x HCX PL APO oil lens and standard acquisition software and detectors were used. Once acquired Pearson’s correlation colocalization and signal intensity analyses were quantified using Volocity 6.3 software (PerkinElmer). Image and line scan analysis was completed using ImageJ FIJI software. GraphPad Prism 7 was used for presentation and statistical analysis of data.

Live cell imaging was performed using a Leica SP8 AOBS confocal laser scanning microscope attached to a Leica DM I6000 inverted epifluorescence microscope. The adaptive focus control was used to prevent drift of the Z plane over time. The two hybrid GaAsP detectors were used to enable low laser settings. Images were acquired using the 63x HC PL APO CS2 lens and a speed of one image per 10 seconds. Imaging was performed at 37°C and 2x rapalog DMEM complete media was added to the pre-selected cell.

## ACKNOWLEDGEMENTS

This work was funded through Wellcome Trust (104568/Z/14/Z) and the Medical Research Council (MR/L007363/1 and MR/P018807/1) awards to P.J.C.

## AUTHOR CONTRIBUTIONS

Conceptualization: A.J.E. and P.J.C.; reagent generation: A.J.E., J.L.D. and B.S.; acquisition and analysis of data: A.J.E., J.L.D. and P.J.C.; writing (review and editing): all authors; funding acquisition, resources and supervision: P.J.C.

## Supplementary Figures

Supplementary Movies 1 - **Temporal resolution of retromer knocksideways onto acceptor compartments after the addition of rapalog.** (A) Retromer (peroxisomal acceptor compartment) knocksideways HeLa cells were imaged with a frame being taken every 10 seconds after the addition of rapalog. The merged panel shows both the VPS35-GFP-FRB and PEX-RFP-FKBP channels. Scale bars = 10 µm. Time display 00:00 min:sec. (B) Retromer (mitochondrial acceptor compartment) knocksideways HeLa cells were imaged with a frame being taken every 10 seconds after the addition of rapalog. The merged panel shows both the VPS35-GFP-FRB and Mito-RFP-FKBP channels. Scale bars = 10 µm. Time display 00:00 min:sec.

**Supplementary Figure 1.**
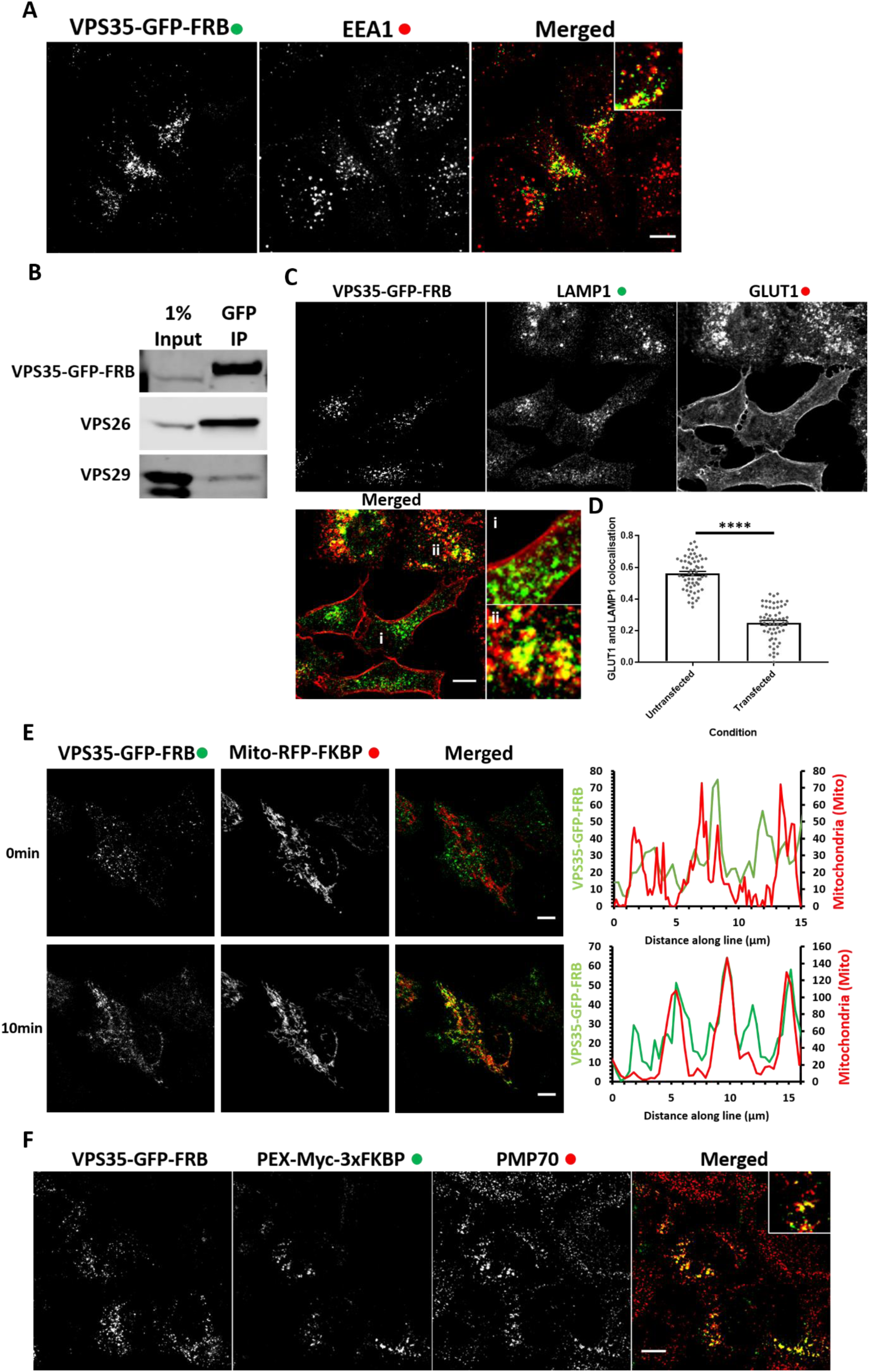
Retromer knocksideways is dynamic and is functionally active. (A) VPS35 knocksideways cells were fixed and then labelled with anti-EEA1. Both the VPS35-GFP-FRB and EEA1 channels are shown in the merged panel with a zoom in panel. Scale bars = 10 µm (B) Western blot analysis of a GFP immunoprecipitation of VPS35-GFP-FRB using anti-VPS35, anti-VPS26 and anti-VPS29. (C) VPS35 knockout cells were transfected with retromer knocksideways and then fixed and labelled with anti-LAMP1 and anti-GLUT1. The merged panel shows both LAMP1 and GLUT1 channels and two zoom-in panels are displayed showing a VPS35-GFP-FRB expressing cell (i) and a cell not expressing retromer (ii, VPS35 knockout cell). Scale bars = 10 µm. (D) Pearsons co-localization between LAMP1 and GLUT1 in untransfected and transfected cells, *n*_exp_ = 3, *n*_cell_ = 60 with all data points being displayed. Statistical analysis performed – Welch’s t-test ****<0.0001. (E) HeLa cells transfected with retromer knocksideways (VPS35-GFP-FRB and Mito-RFP-FKBP). Still frames are shown from a movie (Supplementary Movies 1B) at either 0 minutes or 10 minutes after the addition of rapalog. Line scans were generated using ImageJ drawing a line through mitochondrial structures and represent the colocalization between VPS35-GFP-FRB and Mito-RFP-FKBP at each timepoint. The merged panel displays both channels. Scale bars = 10 µm. (F) HeLa cells were transfected with retromer knocksideways (Peroxisomal, PEX-myc-3xFKBP acceptor compartment) and then antibody labelled with anti-myc and the peroxisomal marker anti-PMP70. The merged panel displays both PEX-myc-3xFKBP and PMP70. Scale bars = 10 µm.

**Supplementary Figure 2.**
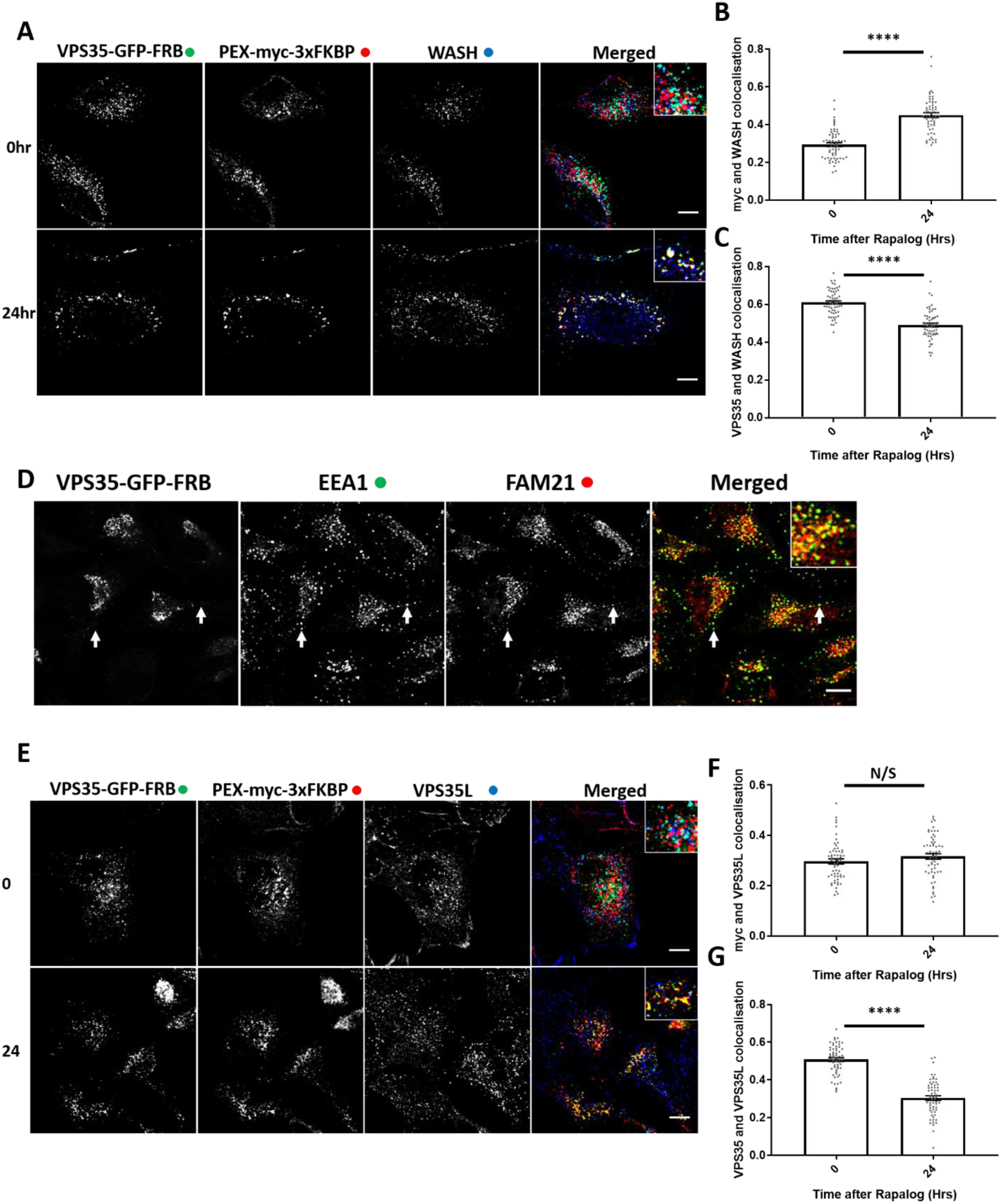
Retromer knocksideways used as an *in cellulo* interaction assay. (A) Retromer knocksideways HeLa cells were fixed before or after 24 hours of rapalog addition and then labelled with anti-myc and anti-WASH. The merged panel displays all three channels with zoom-in panels. Scale bars = 10 µm. (B) Pearsons colocalization between myc and WASH before and after 24 hours of rapalog, *n*_exp_ = 3, *n*_cell_ = 60 with all data points being displayed. Statistical analysis performed – Welch’s t-test, ****<0.0001. (C) Pearsons co-localization between VPS35-GFP-FRB and WASH before and after 24 hours of rapalog, *n*_exp_ = 3, *n*_cell_ = 60 with all data points being displayed. Statistical analysis performed Welch’s t-test ****<0.0001. (D) Retromer knocksideways HeLa were fixed after 24 hours of rapalog addition and stained using anti-EEA1 and anti-FAM21. The merged panel displays both the EEA1 and FAM21 channels with a zoom in panel. Scale bars = 10 µm. Arrows on each panel display areas of co-localization between EEA1 and FAM21. (E) Retromer knocksideways HeLa cells were fixed before or after 24 hours of rapalog addition and then labelled with anti-myc and anti-VPS35L. The merged panel displays all three channels with zoom-in panels. Scale bars = 10 µm. (F) Pearsons co-localization between myc and VPS35L before and after 24 hours of rapalog, *n*_exp_ = 3, *n*_cell_ = 60 with all data points being displayed. Statistical analysis performed - Welch’s t-test, N/S >0.05. (G) Pearsons colocalization between VPS35-GFP-FRB and VPS35L before and after 24 hours of rapalog, *n*_exp_ = 3, *n*_cell_ = 60 with all data points being displayed. Statistical analysis performed – Welch’s t-test, ****<0.0001.

**Supplementary Figure 3.**
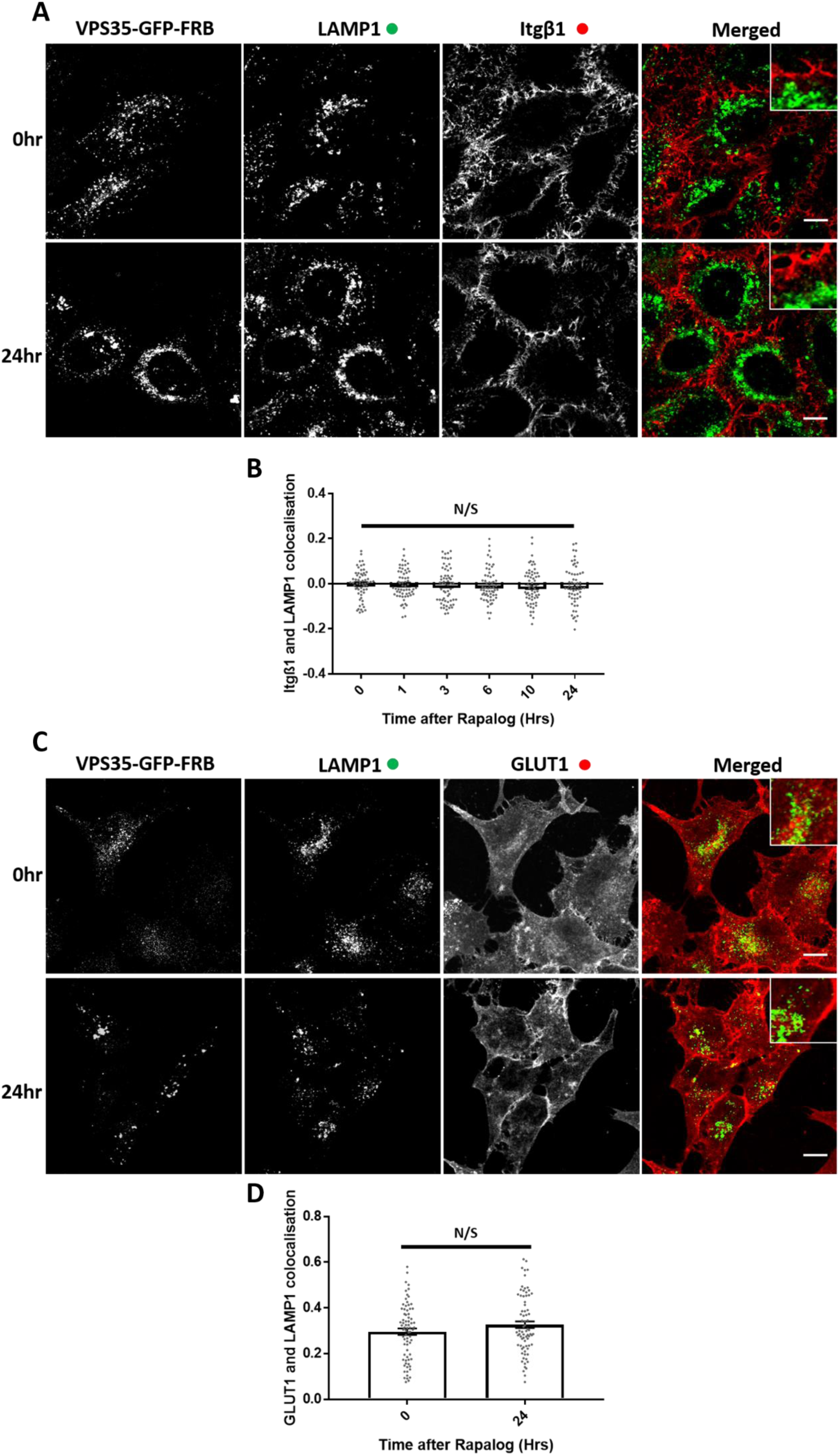
Retromer knocksideways specifically inactivates retromer recycling. (A) Retromer knocksideways HeLa cells were fixed before and after the addition of rapalog for multiple timepoints and then labelled for anti-LAMP1 and anti-Itgβ1. The merged panel shows both the LAMP1 and Itgβ1 channels with a zoom panel. Scale bars = 10 µm. (B) Pearsons co-localization between LAMP1 and Itgβ1 before and after multiple timepoints of rapalog addition, *n*_exp_ = 3, *n*_cell_ = 60 with all data points being displayed. Statistical analysis performed - Ordinary one-way ANOVA with multiple comparisons, N/S >0.05. (C) Retromer knocksideways was transfected into wild-type HeLa cells and were fixed before and after the addition of rapalog for 24 hours and then labelled for anti-LAMP1 and anti-GLUT1. The merged panel shows both the LAMP1 and GLUT1 channels with a zoom panel. Scale bars = 10 µm. (D) Pearsons co-localization between LAMP1 and GLUT1 before and after 24 hours of rapalog, *n*_exp_ = 3, *n*_cell_ = 60 with all data points being displayed. Statistical analysis performed – Welch’s t-test, N/S >0.05.

**Supplementary Figure 4.**
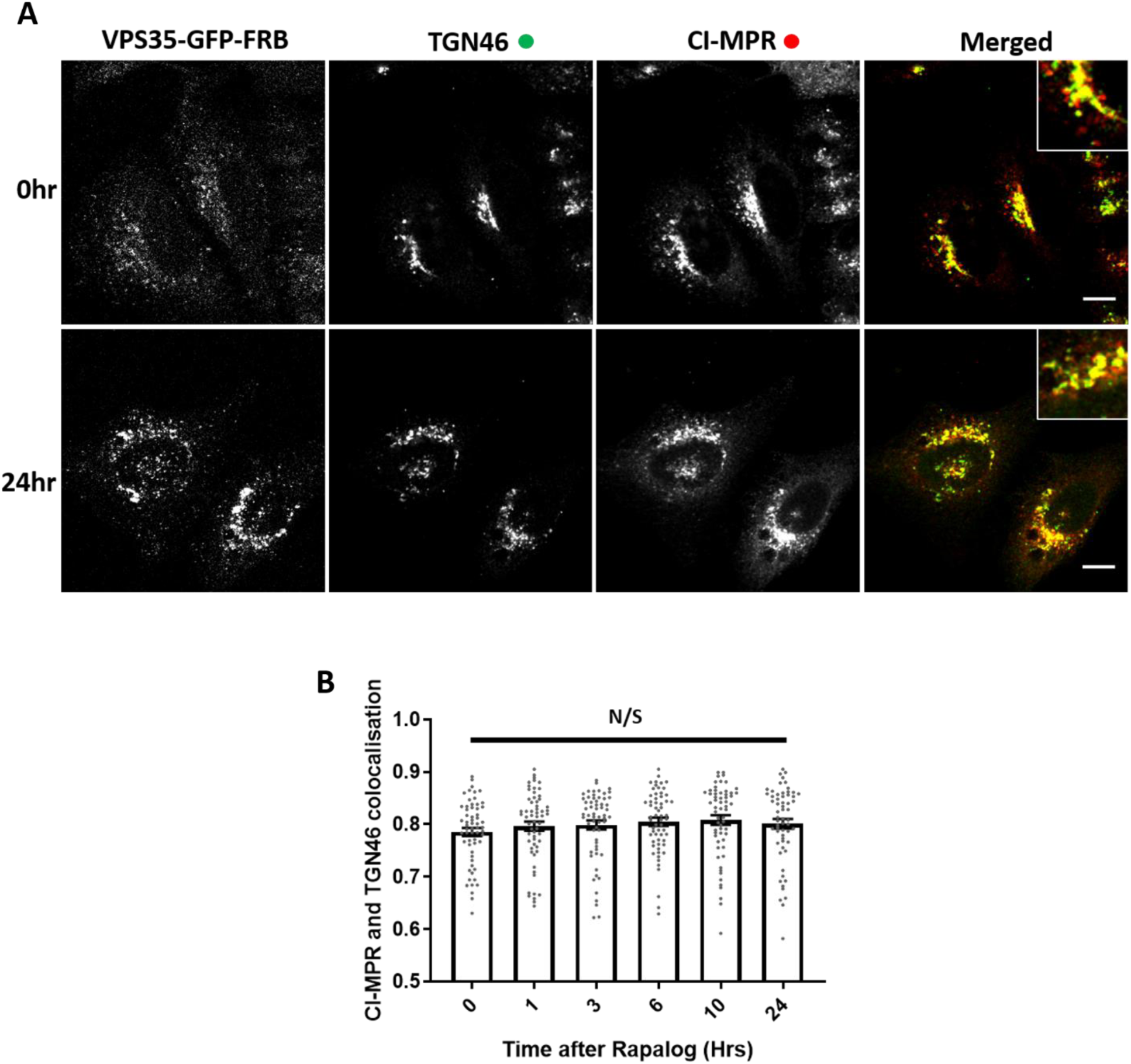
Knocksideways reveals not detectable affect of retromer in retrograde CI-MPR transport. (A) Retromer knocksideways HeLa cells were fixed before or after multiple timepoints of rapalog addition and labelled for anti-TGN46 and anti-CI-MPR. A merged panel displays both the TGN46 and CI-MPR channels and has a zoom panel. Scale bars = 10 µm. (B) Pearsons co-localization between CI-MPR and TGN46 at multiple timepoints, *n*_exp_ = 3, *n*_cell_ = 60 with all datapoints being displayed. Statistical analysis performed - Ordinary one-way ANOVA with multiple comparisons, N/S >0.05.

**Supplementary Figure 5.**
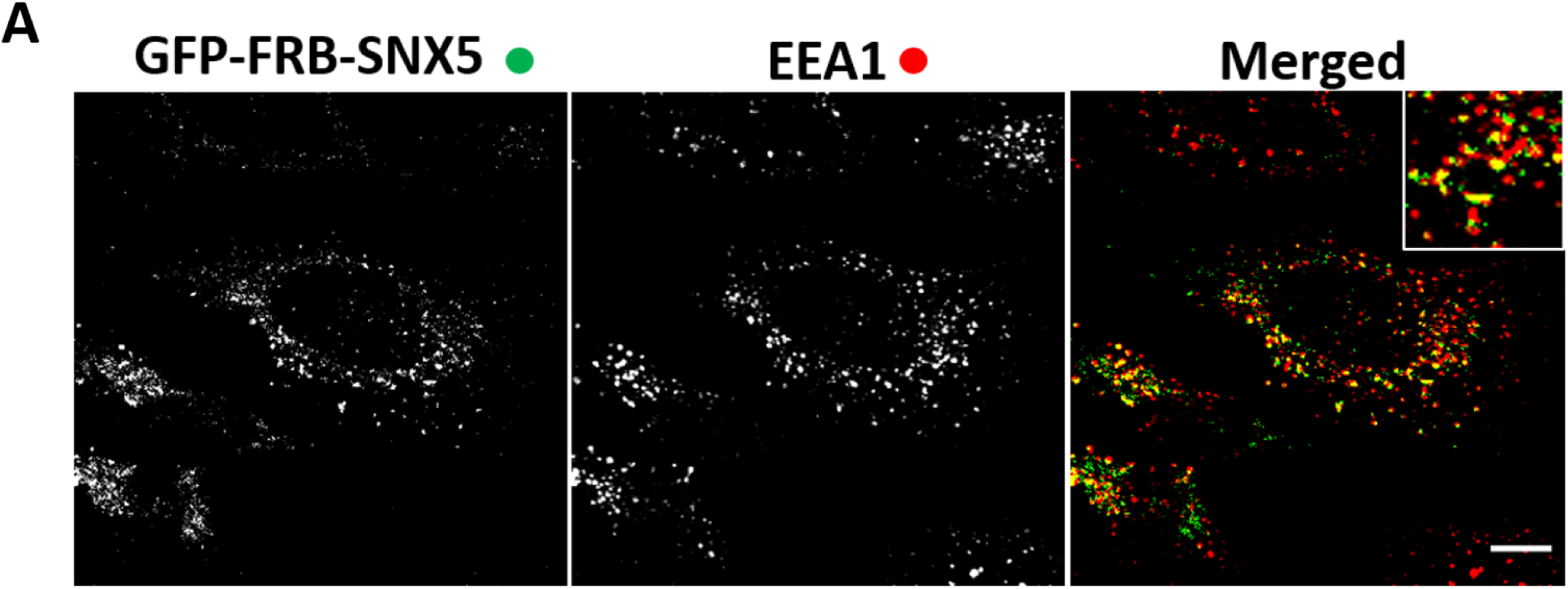
ESCPE-1 knocksideways drags endosomal and peroxisomal compartments together and removes SNX-BARs from endosomes. (A) SNX-BAR knocksideways cells were fixed and then labelled with anti-EEA1. Both the VPS35-GFP-FRB and EEA1 channels are shown in the merged panel with a zoom in panel. Scale bars = 10 µm.

**Supplementary Figure 6.**
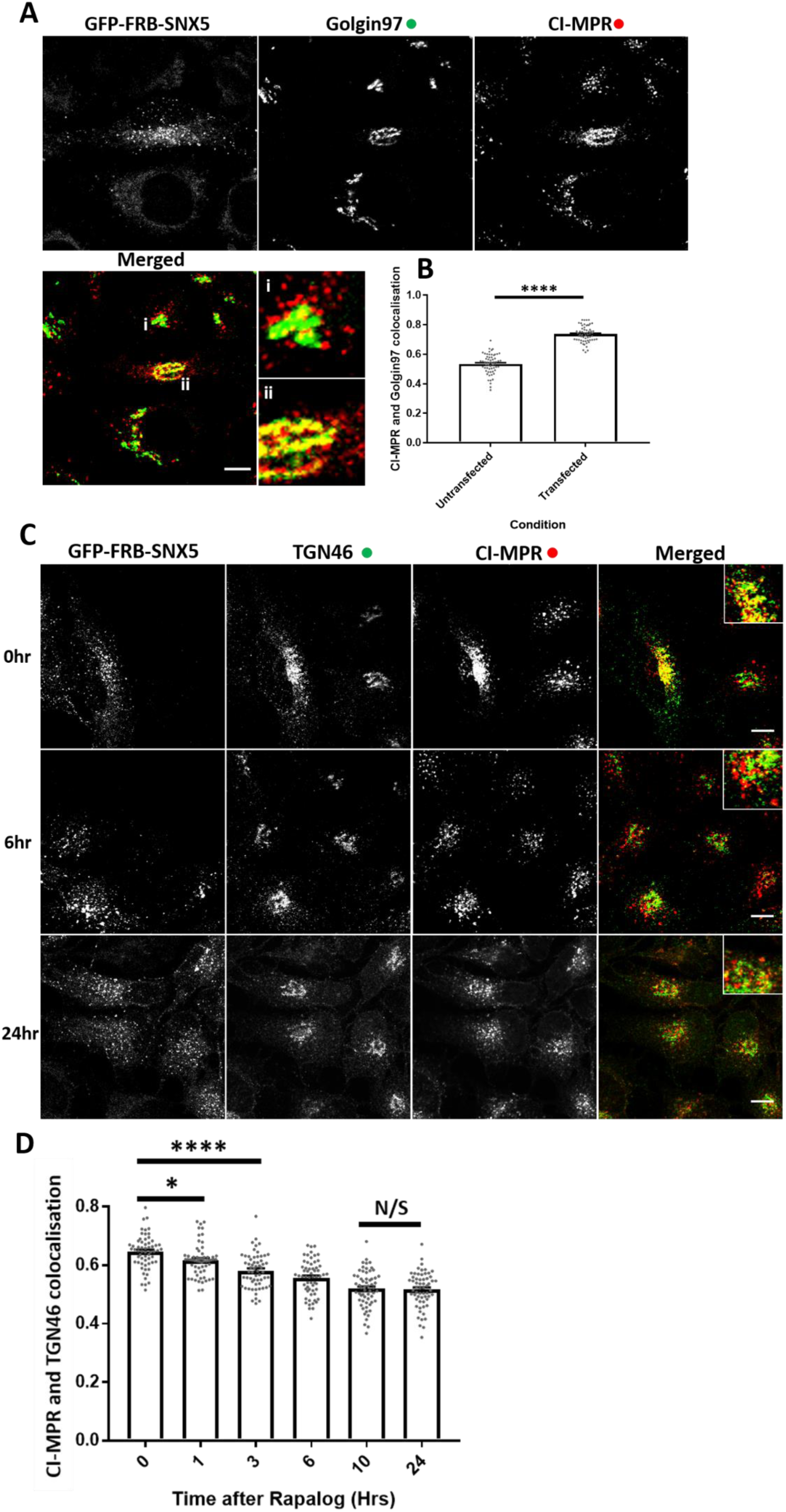
ESCPE-1 knocksideways rescues SNX5/SNX6 dual knockout cells phenotypes and allows the time-resolved redistribution of CI-MPR redistribution after rapalog. (A) Dual SNX5/SNX6 knockout cells were transfected with SNX-BAR knocksideways and fixed and labelled with anti-Golgin97 and anti-CI-MPR. The merged panel shows Golgin97 and CI-MPR channels and two zooms showing a cell not expressing GFP-FRB-SNX5 (i, SNX5/SNX6 knockout cell) and a transfected GFP-FRB-SNX5 expressing cell (ii). Scale bars = 10 µm. (B) Pearsons co-localization between Golgin97 and CI-MPR in untransfected and transfected cells, *n*_exp_ = 3, *n*_cell_ = 60 with all data points being displayed. Statistical analysis performed – Welch’s t-test, ****<0.0001. (B) SNX-BAR knocksideways HeLa cells were fixed before and after the addition of rapalog and labelled for anti-TGN46 and anti-CI-MPR. The merged panel shows both the TGN46 and CI-MPR channels with a zoom panel. Scale bars = 10 µm. (D) Pearsons co-localization between TGN46 and CI-MPR before and after multiple timepoints of rapalog, *n*_exp_ = 3, *n*_cell_ = 60 with all data points being displayed. Statistical analysis performed - Ordinary one-way ANOVA with multiple comparisons, ****<0.0001, *<0.05, N/S >0.05.

**Supplementary Figure 7.**
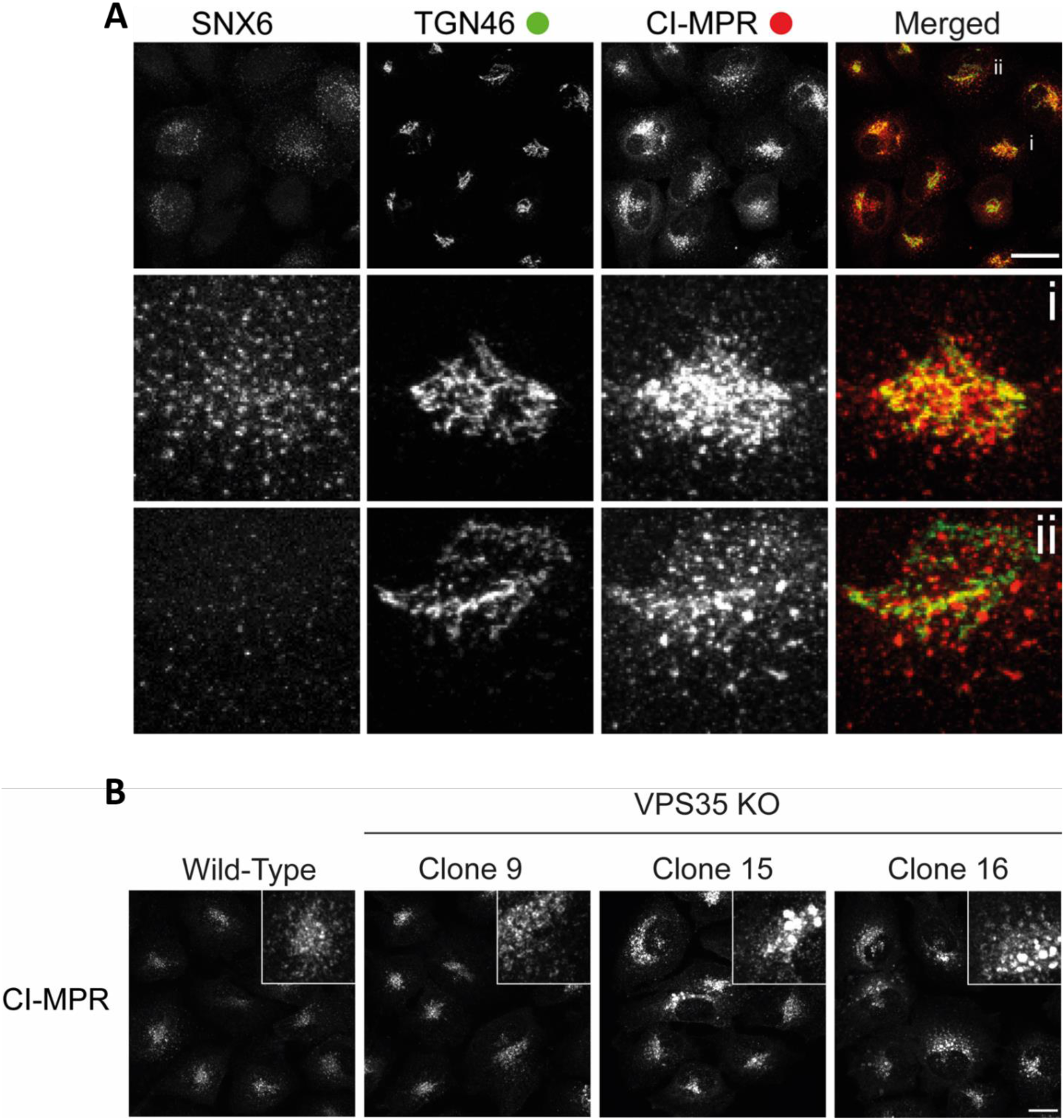
CI-MPR expression is upregulated in VPS35 knockout H4 cells and redistributed away from the *trans*-Golgi network in SNX5/SNX6 knockout H4 cells. (A) Dual SNX5/SNX6 knockout H4 neuroblastoma cells mixed population were fixed and stained with anti-CI-MPR, anti-TGN46 and anti-SNX6. The merged panel shows both TGN46 and CI-MPR and two zooms showing a (i) SNX5/SNX6 positive cell and a (ii) SNX5/SNX6 knockout cell. Scale bars = 20 µm. (B) VPS35 knockout clonal cells fixed and stained with anti-CI-MPR. Scale bars = 20 µm.

**Supplementary Figure 8.**
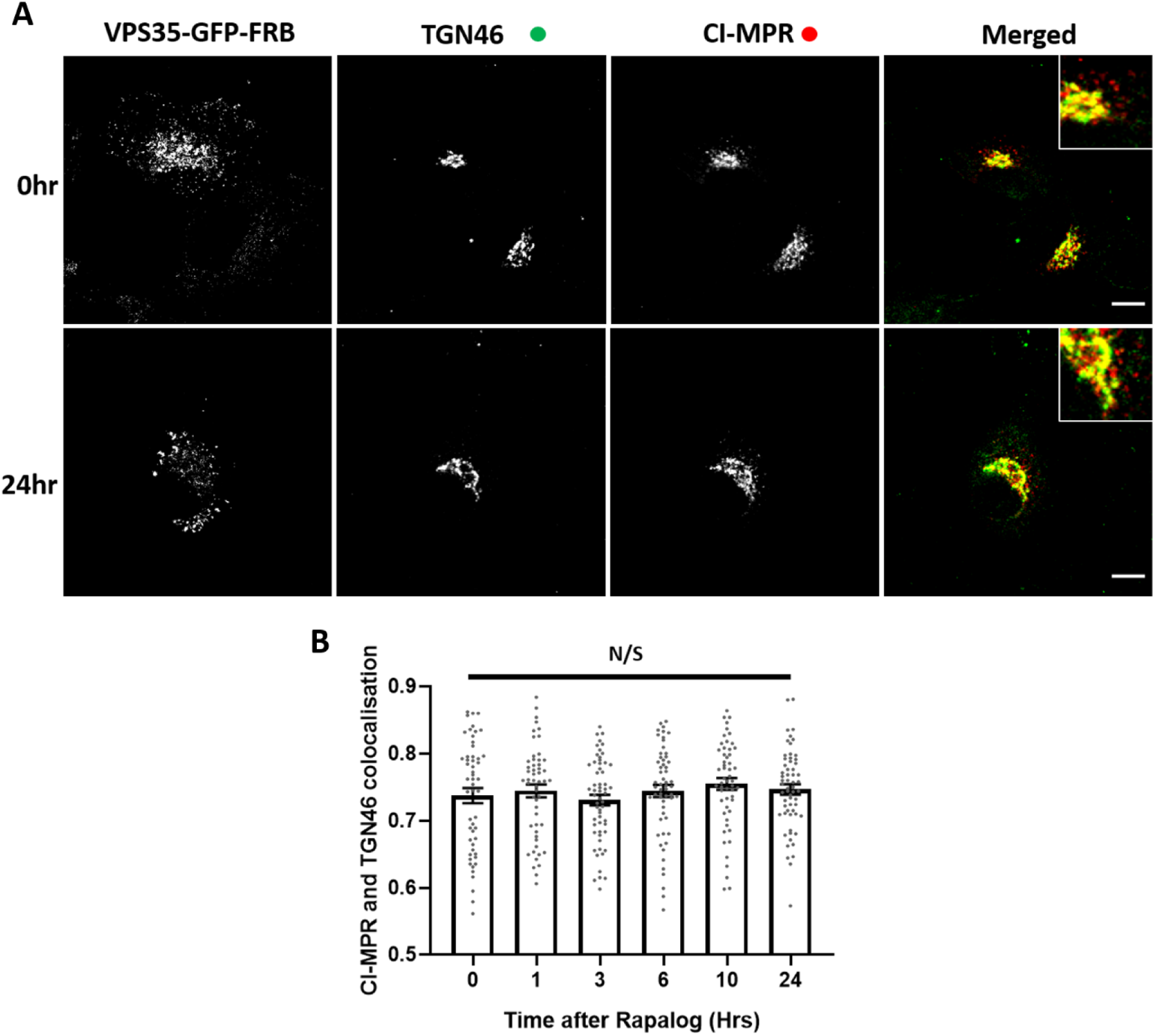
VPS35 knocksideways in H4 neuroblastoma cells demonstrate no visible role for retromer in the retrograde trafficking of CI-MPR. (A) Retromer knocksideways H4 neuroblastoma cells were fixed before or after multiple time points of rapalog addition and labelled with anti-TGN46 and anti-CI-MPR. A merged panel displays both the TGN46 and CI-MPR channels. Scale bars = 10 µm. (B) Pearsons co-localization between CI-MPR and TGN46 at multiple timepoints, *n*_exp_ = 3, *n*_cell_ = 52-56 with all datapoints being displayed. Statistical analysis performed - Ordinary one-way ANOVA with multiple comparisons, N/S >0.05.

